# Pyridoxine supplementation confers protection against *SGPL1^R222Q^* variant sphingosine phosphate lyase insufficiency syndrome

**DOI:** 10.64898/2026.05.11.724358

**Authors:** Ranjha Khan, Maria L. Allende, Ehtesham Khalid, Joanna Y. Lee, Everett Stone, Max Rodnick-Smith, Audrey Izuhara, Vadym Buncha, Georgina Gyarmati, Janos Peti-Peterdi, Ranya Al-Khaledy, Jeffrey B. Hodgin, Gizachew Tassew, Babak Oskouian, Rachel Zhang, Richard L. Proia, Julie D. Saba

## Abstract

Sphingosine-1-phosphate lyase insufficiency syndrome (SPLIS) is a rare condition causing nephrotic syndrome, neuropathy, and other manifestations. SPLIS is caused by mutations in *SGPL1*, which encodes sphingosine-1-phosphate lyase (SPL), a pyridoxal 5’-phosphate (PLP)-dependent enzyme needed to degrade the bioactive sphingolipid sphingosine-1-phosphate (S1P). Supplementation with the PLP precursor pyridoxine benefits some individuals with PLP-dependent enzymopathies. We sought to establish whether pyridoxine has therapeutic activity in SPLIS. Neurological improvement, plasma S1P normalization, and increased SPL activity in patient-derived fibroblasts were observed after pyridoxine supplementation in a patient with R222Q-variant SPLIS. Additionally, PLP dose-dependently augmented recombinant R222Q-variant SPL activity. To further explore pyridoxine’s effects, gene editing was employed to create an R222Q-variant SPLIS mouse model. SPL^R222Q^ mice fed pyridoxine-enriched chow lacked obvious phenotypes. However, SPL inactivation, S1P accumulation, wasting, anemia, proteinuria, and glomerulosclerosis developed in SPL^R222Q^ but not WT mice fed chow with reduced pyridoxine. Ultrastructural analysis and super-resolution microscopy showed podocyte loss and foot process effacement. Transcriptional profiling revealed a pattern of cytokine upregulation and extracellular matrix remodeling. Inhibiting S1P production prevented nephrosis in SPL^R222Q^ mice fed chow lacking pyridoxine. Our findings establish a novel SPLIS mouse model that recapitulates R222Q-variant SPLIS, demonstrates its responsiveness to pyridoxine, and implicates S1P in its pathophysiology.

## Introduction

Sphingosine phosphate lyase insufficiency syndrome (SPLIS), also known as nephrotic syndrome type 14 (OMIM**#** 617575), is an inborn error of sphingolipid metabolism (1, 2). SPLIS has an estimated prevalence of 11,000 worldwide (3). The main features of the condition include steroid-resistant nephrotic syndrome (SRNS), adrenal insufficiency, peripheral neuropathy, anemia, and immunodeficiency, which may occur together or independently. When diagnosed in infancy with kidney involvement, SPLIS is associated with an 80% mortality within a few years (4). Currently, treatment is supportive, and there is no targeted therapy for SPLIS.

SPLIS is caused by biallelic inactivating mutations in *SGPL1* (OMIM#603729; HGNC:10817). More than 45 SPLIS-associated *SGPL1* variants have been identified, the most common being the R222Q substitution, which accounts for ∼20% of SPLIS cases (5). *SGPL1* encodes sphingosine phosphate lyase (SPL), an essential, pyridoxal 5’-phosphate (PLP)-dependent enzyme that catalyzes the irreversible cleavage of the bioactive signaling molecule sphingosine-1-phosphate (S1P) in the last step of sphingolipid metabolism (6, 7). Complex biochemical consequences of SPL insufficiency contribute to the pathogenesis of SPLIS. The inability to cleave S1P results in the accumulation of S1P in cells, blood, and tissues (8). This may induce aberrant S1P signaling through the five known S1P receptors, which regulate signaling pathways controlling vascular permeability, leukocyte trafficking, neurotransmission, and pro-inflammatory and pro-fibrotic cytokine signaling (9). Reported receptor-independent effects of S1P on calcium homeostasis, epigenetic gene transcription, cholesterol handling, and mitochondrial function may additionally occur in SPLIS (10, 11). All sphingolipids must eventually be metabolized through the common sphingolipid degradative pathway ending in SPL. Consequently, SPL inactivation leads to the accumulation of upstream sphingolipid metabolites, such as ceramides, which can be cytotoxic, as well as a deficiency of the metabolic products, hexadecenal and ethanolamine phosphate, which have their own cellular functions, such as in autophagy and phospholipid biosynthesis (12–16). An ideal SPLIS treatment would address all the biochemical consequences of SPL insufficiency. This could be accomplished by either augmenting endogenous SPL activity or providing a working copy of the *SGPL1* gene or SPL protein, thereby restoring sphingolipid metabolism to the key tissues that need it.

PLP is the active form of vitamin B6 and one of six B6 “vitamers”, the other five forms being enzymatically convertible to PLP *in vivo* (17). Clinical guidelines for certain infantile seizure disorders began recommending the use of the B6 vitamer pyridoxine in the 1980s, and its use in pediatrics is well established. Pyridoxine also has therapeutic activity in some inborn errors of metabolism involving PLP-dependent enzymes (18). Two mechanisms of action have been implicated. By increasing intracellular PLP concentrations, pyridoxine supplementation may overcome the poor cofactor-binding affinity of some pathogenic enzyme variants. Alternatively, PLP may act as a chaperone, stabilizing variants that are prone to misfolding (19). These two mechanisms are not mutually exclusive.

We hypothesized that supplementation with pyridoxine would benefit a subset of SPLIS patients harboring pyridoxine-responsive SPL variants. We had previously shown that pyridoxine supplementation led to clinical and biochemical improvements in two SPLIS patients, including one homozygous for the SPL-R222Q variant (20). As there are currently no targeted treatments for SPLIS and, given the favorable safety profile of pyridoxine and its widespread availability, establishing the efficacy of pyridoxine supplementation in SPLIS (or a tractable model of SPLIS) and identifying pyridoxine-responsive SPL variants are clinically valuable goals. However, the genetic and phenotypic heterogeneity of SPLIS, its rarity, and the difficulty of convincing patients to participate in an interventional study when the supplement is freely available make conducting a pyridoxine randomized clinical trial in SPLIS extremely challenging.

In this study, we aimed to determine whether pyridoxine supplementation can prevent disease progression and reverse organ damage in SPLIS using a combination of preclinical approaches, including the use of patient-derived skin fibroblasts, purified recombinant SPL protein, and a novel, personalized mouse model of R222Q-variant SPLIS. Our results establish SPL^R222Q^ mice as a new, inducible animal model of SPLIS that exhibits a spectrum of SPLIS phenotypes when vitamin B6 is limiting and that are prevented by pyridoxine enrichment. Our cumulative results demonstrate the efficacy of pyridoxine in the treatment of SPLIS caused by the most common pathogenic variant of SPL, the R222Q substitution. Further, we show that preventing S1P production *in vivo* using sphingosine kinase inhibitors protects against the development of SPLIS nephrosis, implicating the accumulation of S1P as the culprit in the pathophysiology of the condition.

## Results

### Pyridoxine attenuates disease manifestations in a patient with SPL R222Q-variant SPLIS

An 18-year-old female presented with peripheral numbness, burning sensation, and pain in the upper and lower extremities that had begun at 8 years of age with periodic improvement over time. Family history was significant for parental consanguinity, two miscarriages, and the death of two female siblings. One sibling had delayed milestones, skin hyperpigmentation, and died at two and a half years of age due to a fever. The other died at two years of age with kidney problems and proteinuria (**Figure S1A**). The patient’s baseline nerve conduction study (NCS) was consistent with axonal polyneuropathy with no evidence of demyelination. Whole-exome sequence (WES) analysis of the proband identified a homozygous *SGPL1* R222Q pathogenic variant previously associated with SPLIS (**Figure S1B**). Both parents were heterozygous for the R222Q variant. The substituted arginine is highly conserved in the eutherian mammals (**Figure S1C**). We previously showed disease modifying activity of cofactor supplementation using the B6 vitamer pyridoxine in two SPLIS subjects, one of whom was homozygous for the R222Q variant (20). To assess whether pyridoxine might be beneficial in the patient, SPL expression was measured by immunoblotting (IB) in whole cell extracts of skin fibroblasts derived from the patient versus a healthy control, both of which were cultured for 96h in medium lacking pyridoxine. A strong SPL protein band was observed in the control fibroblasts, but only a faint band was observed in the SPLIS fibroblasts, whereas SPLIS fibroblasts that were subsequently treated with 100 µM pyridoxine for 72h showed increased signal, indicating a stabilizing effect of cofactor supplementation (**Figure S1D**). Corresponding with these results, baseline SPL enzyme activity in the patient’s fibroblasts was 5% that of healthy control levels, consistent with an inborn error of metabolism (**Figure S1E**). Treatment with pyridoxine resulted in an increase in SPL activity to 25% of control fibroblast levels (**Figure S1E**), a level that correlates with prevention of severe SPLIS phenotypes in murine models of SPL insufficiency (21). Based on this result, a trial of 100 mg/day oral pyridoxine was initiated. Over the next few months, the patient reported an 80% improvement in neurological function, including improved strength and resolution of painful neuralgias. Median motor nerve conduction (**Figure S1F**) as well as sensory nerve conduction in the median and ulnar nerves (data not shown) showed 50% improvement compared to baseline NCS. Concomitantly, the patient’s plasma S1P levels, which were significantly elevated over age-and gender-matched control levels before initiation of pyridoxine treatment, fell into the normal range with treatment (**Figure S1G**).

### PLP treatment stabilizes and dose-dependently augments the catalytic activity of recombinant R222Q-variant SPL

To assess whether PLP supplementation influences the activity of mutant forms of SPL, recombinant WT human SPL and SPLIS-associated R222Q and R222W variants were expressed in *E. coli* and purified initially in the presence of excess PLP, as described in Materials and Methods. The affinity chromatography step yielded a prominent band corresponding to the expected molecular weight of human SPL (∼55 kDa), as visualized on the Coomassie-stained gel with both R222Q and R222W variants exhibiting reduced yields and co-purified with many contaminants that were eliminated with ion exchange chromatography (**Figure 1A**). To characterize the enzymatic activities of WT and variant SPL proteins, a time-course reaction was performed using 50 nM of the holoenzyme and 10 µM of its substrate, S1P. The reaction was conducted in the presence of 10 µM supplementary PLP. The R222Q variant showed enhanced activity with PLP supplementation, while the R222W variant showed virtually no activity with or without PLP (**Figure 1B**). Similarly, we compared the activity of R222Q and R222W variants after normalizing with the WT enzyme. Again, the activity of the R222Q variant was augmented with increasing concentration of PLP, whereas the R222W variant was catalytically dead, and the activity of the WT SPL was not significantly improved by PLP supplementation (**Figure 1C**). Thus, PLP dose-dependently recovered the enzyme activity of the R222Q variant, whereas increasing PLP has little to no effect on the WT enzyme or the R222W variant. We attribute the difference in response of the two different substitutions at residue 222 to the destabilizing effect of steric hindrance caused by the tryptophan substitution compared to glutamine.

**Figure 1.**
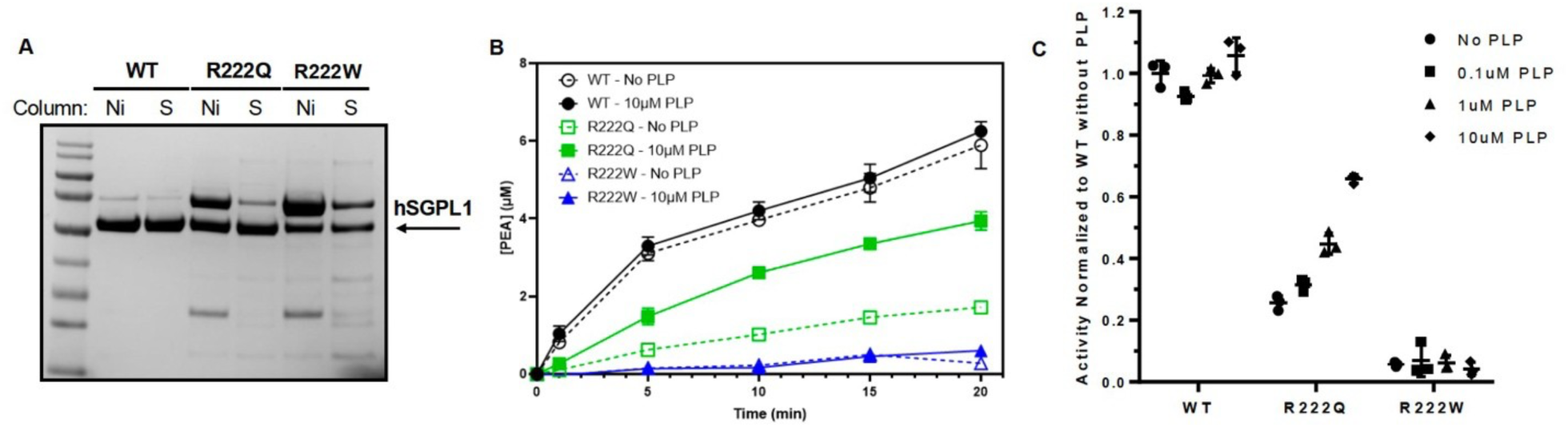
SPL protein purification, stability, and enzymatic activity. (A) Coomassie-stained SDS-PAGE gel of purified Human S1P lyase showing elutions from affinity (Ni) and ion-exchange (S) columns. (B) Time course reaction of 50nM purified S1P lyase with 10 µM S1P with indicated concentrations of PLP. Error bars represent the standard deviation of triplicate measurements. P values, WT 0 uM PLP vs 10 uM = NS, R222Q 0 uM PLP vs 10 uM = 0.00353, and R222W 0 uM PLP vs 10 uM = NS. (C) Activity of enzymes at 5 minutes normalized to the wild-type enzyme without PLP. Enzyme activity with 10uM PLP: WT vs R222Q <0.0001, WT vs R222W <0.0001, and R222Q vs R222W: <0.0001.

### Generation of a murine model of R222Q-variant SPLIS

Based on the above findings suggesting the potential therapeutic activity of pyridoxine in SPLIS, it became important to validate pyridoxine’s prophylactic effects *in vivo*. A well-characterized *Sgpl1* KO mouse line serves as a robust model of SPLIS, characterized by elevated sphingolipids in blood and tissues, SPLIS-associated phenotypes, runting, and death at weaning (7, 22, 23). However, *Sgpl1* KO mice produce no SPL protein, making this an unsuitable preclinical model for testing pyridoxine’s efficacy in SPLIS. Leveraging the fact that mouse and human *SGPL1* gene sequences are identical in the region surrounding the R222 residue, we generated mice harboring the SPL R222Q substitution using CRISPR genome editing technology and verified by DNA sequence, as described in Methods (**Figure 2A**). A single founder female was identified who harbored one *Sgpl1* allele with the intended nucleotide change resulting in a gene product with the R222Q substitution. A novel SphI restriction cleavage site created by the nucleotide change allowed us to detect this variant by PCR followed by DNA digestion with SphI, resulting in two DNA fragments diagnostic of the knock-in mutation (**Figure 2B**). Interestingly, an unintentional *Sgpl1* deletion of 38 base pairs occurred in the *Sgpl1* gene on the founder’s other chromosome (**Figure 2A**). Both the knock-in and deletion mutations in the founder mouse’s DNA were verified by Sanger sequence analysis (**Figure 2A**). Unexpectedly, the founder mouse that harbored knock-in and deletion *Sgpl1* mutations was healthy and reproductive (**Figure 2C**). When bred with a WT mate, she bore pups, some of which carried the R222Q variant. The heterozygous knock-in pups were bred to homozygosity for the SPL R222Q variant, and homozygotes were used for further analysis.

**Figure 2.**
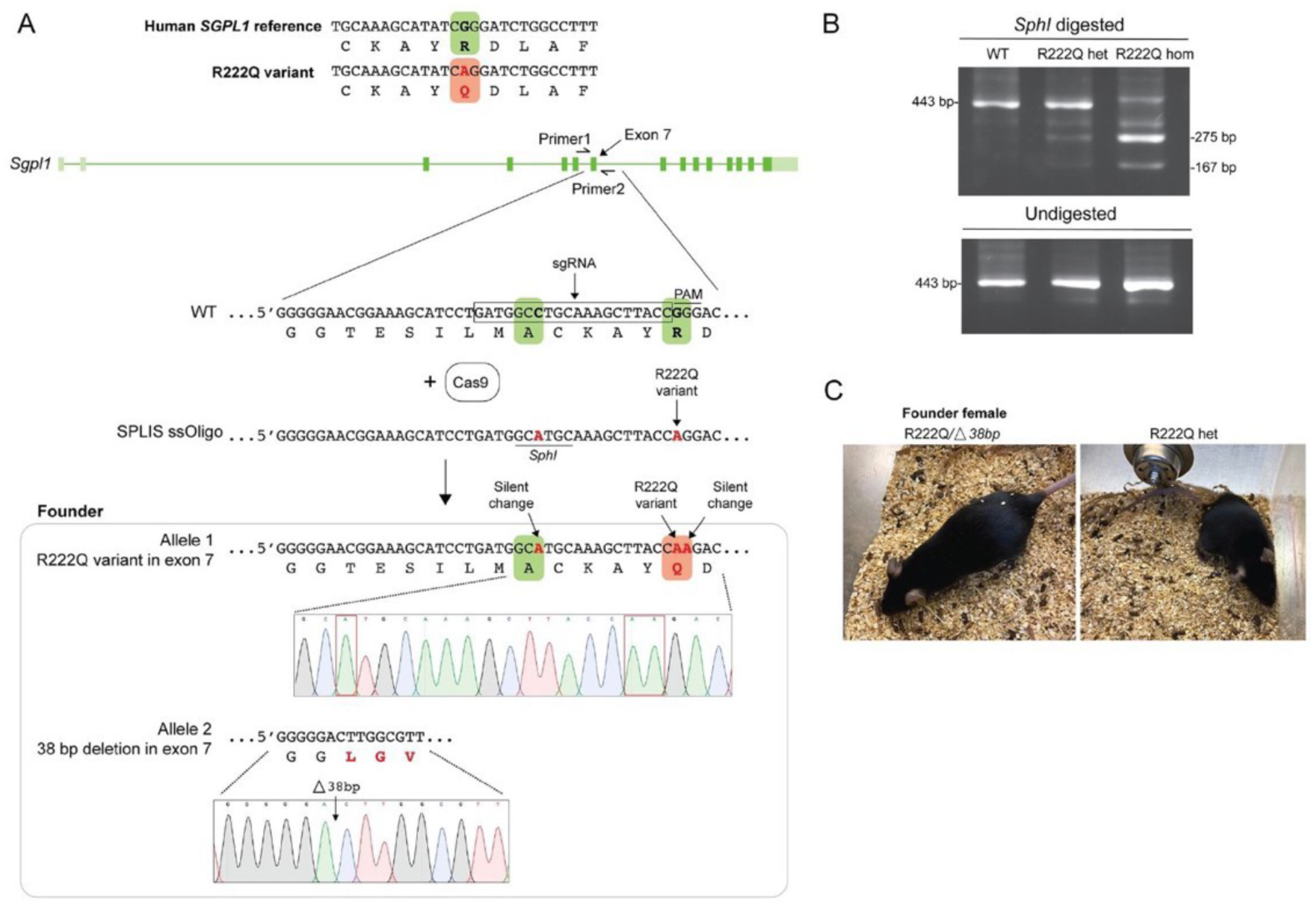
Generation of SPLIS R222Q mutant mice. (A) A single-guide RNA (sgRNA) containing a 20 bp target sequence, 5’ GATGGCCTGCAAAGCTTACC 3’ (Synthego) of exon 7. To introduce the mutant sequence, we designed a single-stranded 199-base oligodeoxynucleotide (SPLIS ssOligo) containing the G-to-A SPLIS mutation and a silent point mutation to create a SphI restriction enzyme site for screening: 5’AGGCTGTACCTCCTGATACCCATCCTTAACTTACTCTGGTTTCTTTTCTTCACATAGGTGACTTCTGGGGGAACGGAAAGCATCCTGATGGCATGCAAAGCTTACCAGGACTTGGCGTTAGAGAAGGGGATCAAAACTCCAGAAATGTATGGATGTGTGTGTTTGTTTCCCTTCTGATATTGTCTATTTGTGGCAGCAC 3’ (Ultramer DNA, IDT). The C57BL/6J strain was used as an embryo donor. Fertilized oocytes were injected with Cas9 protein (TrueCut Cas9 protein V2, Thermofisher), the sgRNA, and the SPLIS ssOligo and transferred to the oviducts of pseudopregnant female mice. (B) Mice were genotyped by PCR. The resulting PCR product (443 bp) was digested with SphI to detect the mutant sequence. After digestion, the WT allele yielded a 443 bp fragment; the SPLIS-modified allele yielded 275 bp and 167 bp bands. We detected 1 founder whose PCR fragment was digested by SphI (out of 99 offspring screened). Sequences of the undigested PCR fragments were determined after TOPO TA cloning (ThermoFisher) by Sanger sequencing. The founder female carried one allele containing R222Q mutation and SphI restriction site and one allele with a 38-bp deletion in exon 7, resulting in a frameshift and premature stop codon (A, B). (C) When mated with a WT male, the founder produced 4 pups, 2 *Sgpl1* R222Q/WT and 2 *Sgpl1* del/WT, segregating the 2 alleles.

### Pyridoxine supplementation augments tissue SPL activity and prevents S1P accumulation in SPL^R222Q^ mice

We originally hypothesized that mice homozygous for the R222Q variant (hereafter referred to as SPL^R222Q^ mice) would exhibit SPLIS-like phenotypes that would be ameliorated by pyridoxine supplementation. However, the healthy state of the founder mouse, which harbored a deletion in one *Sgpl1* allele and relied on the single R222Q variant for function, showed no obvious phenotype under standard husbandry conditions and given regular mouse chow. Similarly, homozygous SPL^R222Q^ mice showed no obvious phenotype under the same conditions. Maintenance pyridoxine requirements for mice are estimated to be 1 mg/kg/d (24). Based on the pyridoxine content of regular laboratory mouse chow and an estimated intake of 5g chow/day, we calculated that the mice were receiving 6-7 mg/kg/d pyridoxine, i.e., 6 to 7-fold higher than the minimum daily requirements. Thus, standard mouse chow contains supplementation with what could represent a “therapeutic” amount of pyridoxine. To determine the impact of eliminating pyridoxine supplementation on the R222Q variant, we compared the SPL expression in vital organs of SPL^R222Q^ mice given standard mouse chow (Hi-B6) versus chow measuring 2 ppm pyridoxine with No Added pyridoxine (NA-B6) for 2 months. Importantly, to ensure we were not eliciting phenotypes caused by general vitamin B6 deficiency, all experiments also included groups of WT mice on Hi-B6 and NA-B6 chow for comparison. IB of whole kidney extracts revealed robust SPL protein expression in WT kidneys regardless of diet, indicating that the WT SPL abundance is unaffected when pyridoxine supplementation is withdrawn for two months (**Figure 3A**). This is consistent with the lack of effect of increasing amounts of PLP on purified WT SPL (**Figure 1B-C**) and can be explained by the fact that the WT SPL polypeptide folds into its native conformation, is abundant and stable, and exhibits a high affinity for PLP. In contrast, SPL protein levels were reduced in the kidneys of SPL^R222Q^ mice fed Hi-B6 chow, with even lower SPL levels observed in SPL^R222Q^ mice fed NA-B6 chow (**Figure 3A**). Consistent with the WT expression pattern, SPL activity levels in WT mouse liver, lung, kidney, and brain tissues were comparable on either diet (**Figure 3B-E**). SPL activity in the tissues of SPL^R222Q^ mice fed Hi-B6 chow was 25-50% of WT, a level shown to prevent most SPLIS-associated phenotypes in mice and also consistent with the lack of reported disease phenotypes in human SPLIS carriers (21). In contrast, SPL activity in the liver, lung and kidney tissues of SPL^R222Q^ mice fed NA-B6 chow decreased to 1-5% of the levels observed in the tissues of WT mice on the same chow, a range comparable to that of an inborn error of metabolism, with somewhat more residual SPL activity observed in brain (**Figure 3B-E**). S1P levels in the plasma and tissues of WT and SPL^R222Q^ mice fed Hi-B6 chow and WT mice on NA-B6 chow were similar (**Figure 3F-J**). This indicates that the residual global SPL activity in SPL^R222Q^ mice fed Hi-B6 chow is sufficient to maintain normal sphingolipid metabolism. In contrast, plasma and tissue S1P levels of SPL^R222Q^ mice fed NA-B6 chow were highly elevated compared to the control groups (**Figure 3F-J**).

**Figure 3.**
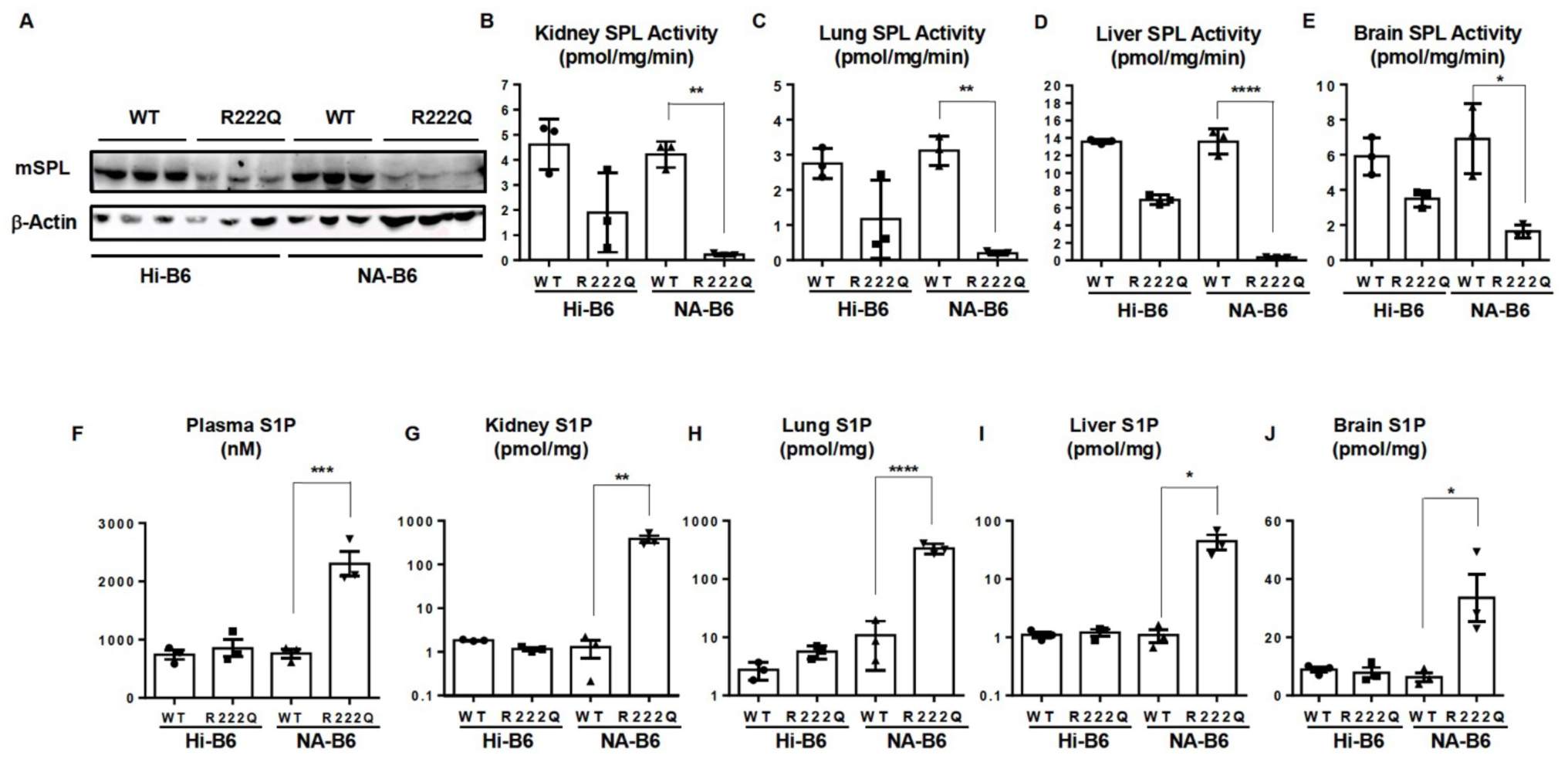
SPL^R222Q^ mice on no added B6 (NA-B6) manifest severe SPL insufficiency and elevated S1P level. (A) Immunoblot from the kidneys of WT, SPL^R222Q,^ with and without added B6, showing reduced level of SPL protein only in R222Q mice with NA-B6 chow. NA-B6 = no added B6 in the chow. (B-E) SPL activity in vital organs (kidney, lung, liver, and brain) of WT and SPL^R222Q^ mice, confirming drastically low activity of SPL in SPL^R222Q^ mice fed NA-B6 chow. (F-J) Measurement of S1P levels in the plasma, kidney, lung, liver, and brain confirms higher levels of S1P in the organs and plasma of SPL^R222Q^ mice fed on NA-B6 chow. * p ≤ 0.05, ** p ≤ 0.01, *** p ≤ 0.001, and **** p ≤ 0.0001.

### Pyridoxine supplementation prevents development of SPLIS phenotypes in SPL^R222Q^ mice

We next investigated whether pyridoxine supplementation suppresses the development of SPLIS-related phenotypes in SPL^R222Q^ mice. Grip strength (**Figure 4A**) and body weight (**Figure 4B**), which represent measures of neurological function and general health, respectively, deteriorated in SPL^R222Q^ mice after two months on NA-B6 chow, compared to all other groups. Urine albumin creatinine ratio (ACR), a measure of proteinuria, rose significantly in SPL^R222Q^ mice after three weeks on NA-B6 chow (**Figure 4C**), and this increased further after two months on NA-B6 chow (**Figure 4D**). Correspondingly, serum albumin levels were lower in SPL^R222Q^ mice after two months on NA-B6 chow compared to all other groups (**Figure 4E**). RBC indices (hemoglobin, hematocrit, and RBC mass) were also lower, indicating the development of anemia in SPL^R222Q^ mice after two months on NA-B6 diet, compared to all other groups. (**Figure 4F-H**).

**Figure 4.**
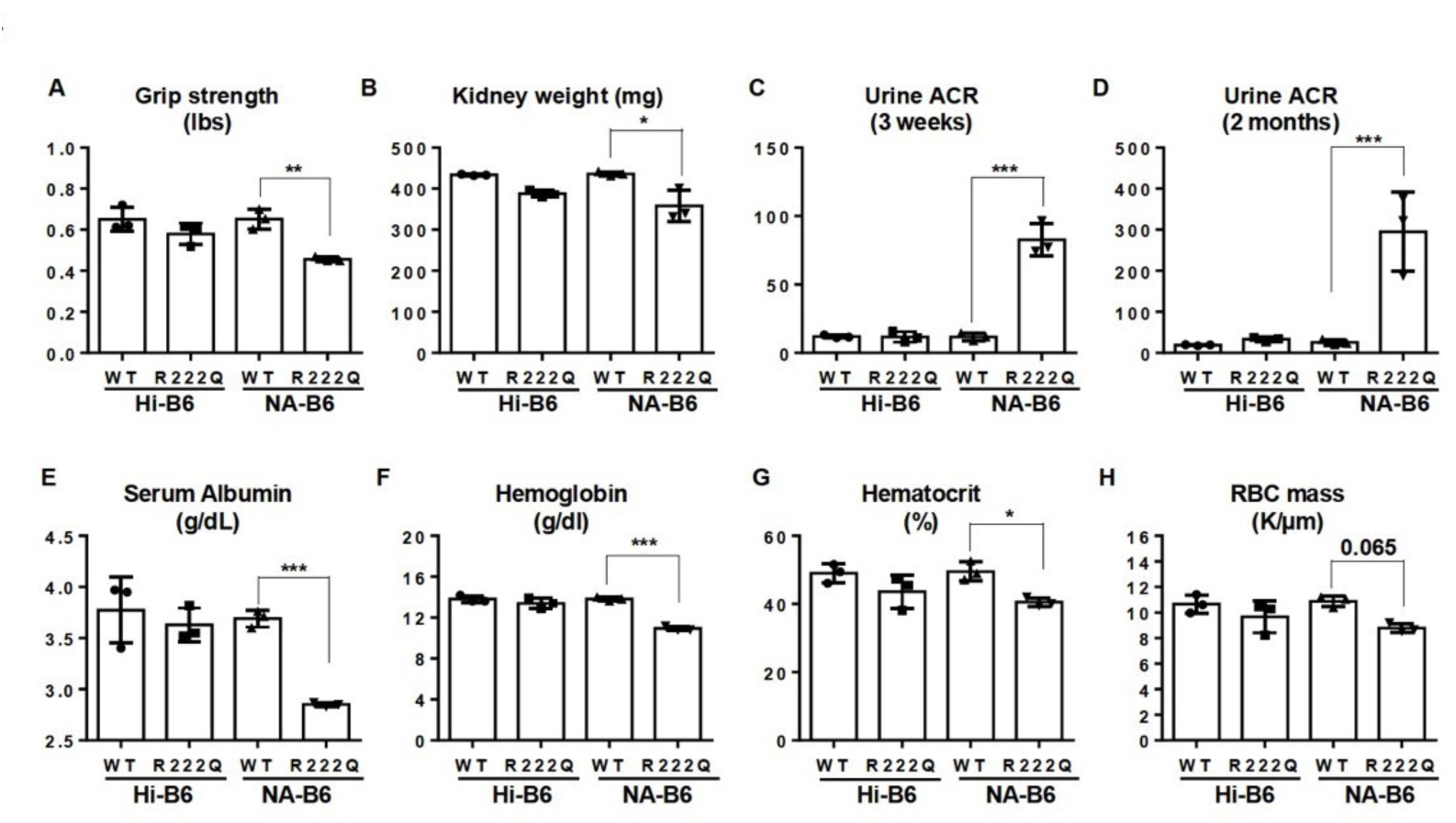
Pyridoxine supplementation ameliorates SPLIS features in SPL^R222Q^ mice. (A) Improvement in grip strength observed with B6 supplementation. (B) Body weight comparison. (C-D) Urine ACR at 3 week and 8-week time points, showing improvement in SPL^R222Q^ mice on Hi-B6 chow. (E) Serum albumin level. (F-H) RBC indices. * p ≤ 0.05, ** p ≤ 0.01, and *** p ≤ 0.001.

### Pyridoxine supplementation prevents glomerulosclerosis, podocyte loss, and foot process effacement in SPL^R222Q^ mice

Histology of periodic acid Schiff (PAS)-stained kidney sections of WT and SPL^R222Q^ mice on Hi-B6 chow and WT mice on NA-B6 chow was normal in appearance and indistinguishable from one another (**Figure 5 A-C**). In contrast, PAS staining of SPL^R222Q^ kidneys after two months on NA-B6 chow showed glomerulosclerosis with collagen deposition, mesangial expansion, and protein casts (**Figure 5D**). Quantification of abnormal glomeruli indicated that only SPL^R222^ mice, after two months on NA-B6 chow, showed 8% focal glomerulosclerosis and 13% global glomerulosclerosis (**Table S1**). Injury and death of glomerular podocytes, which maintain the kidney filtration barrier, are considered the main underlying cause of steroid-resistant nephrotic syndrome (25). Loss of podocyte foot processes indicates podocyte injury and is a pathognomonic sign of nephrotic syndrome. Ultrastructural analysis of the kidney cortices of SPL^R222Q^ mice fed NA-B6 chow by transmission electron microscopy (TEM) revealed podocyte foot process effacement and glomerular basement membrane thickening, features not observed in control kidneys (**Figure 5E-F**). Stimulated emission depletion (STED) super-resolution microscopy of kidney cortices stained with antibodies against the podocyte marker nephrin demonstrated reduced slit diaphragm density (podocyte effacement) and overall reduced nephrin immunolabeling, indicating podocyte drop-out in the kidneys of SPL^R222Q^ mice on NA-B6 chow compared to controls (**Figure 5G-J**).

**Figure 5.**
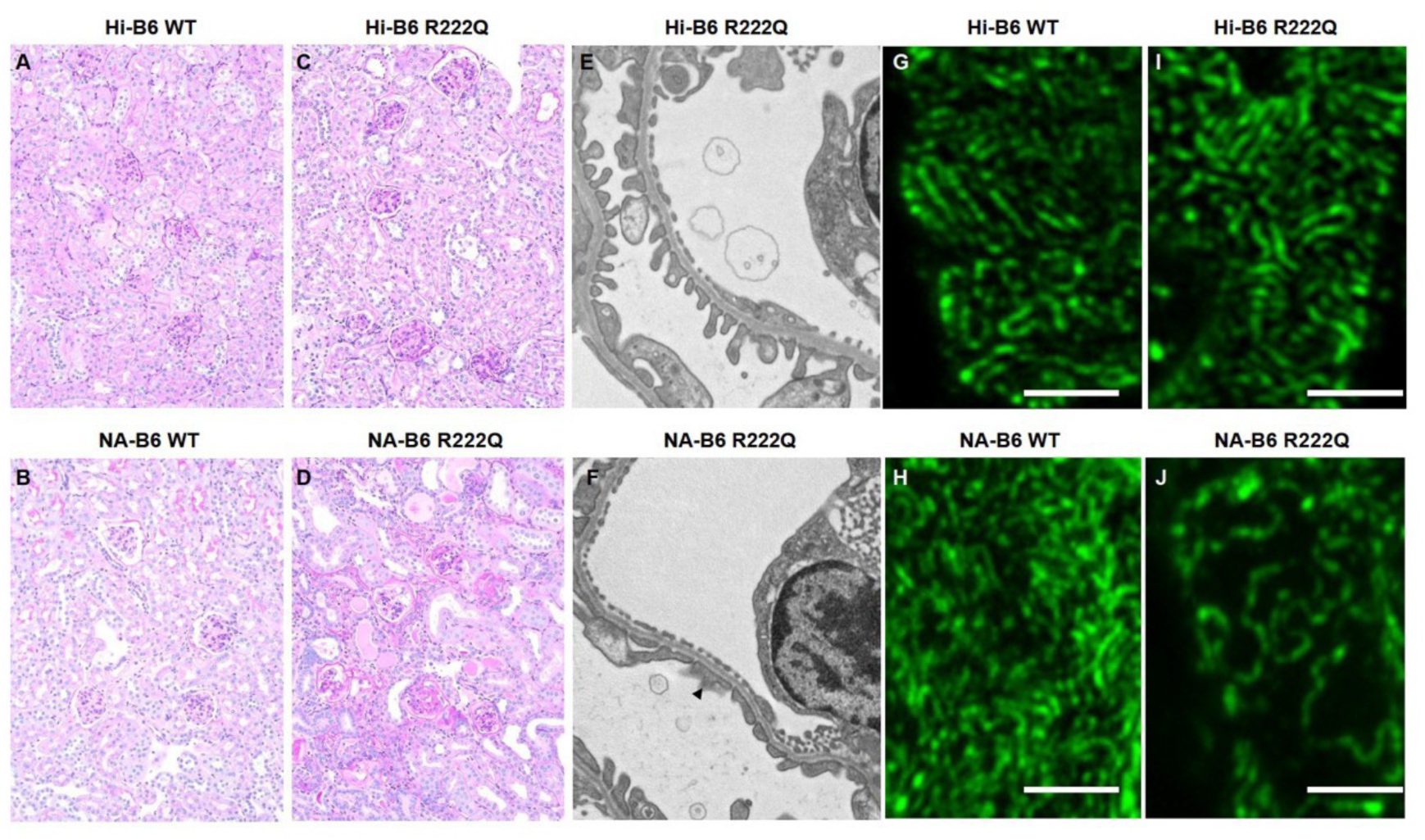
Addition of Hi-B6 in the chow prevents glomerulosclerosis, podocyte loss, and foot process effacement in SPL^R222Q^ mice. (A-D) PAS staining shows glomerulosclerosis in SPL^R222Q^ mice that were fed on NA-B6 chow, while SPL^R222Q^ mice on Hi-B6 and WT mice on either diet exhibited normal morphology of glomeruli and tubules. (E-F) Transmission electron microscopy reveals podocyte effacement in SPL^R222Q^ mice that were fed on NA-B6 chow compared to control mice on NA-B6 chow. (G-H) STED imaging reveals disrupted foot process morphology in SPL^R222Q^ mice fed NA-B6 chow compared to SPL^R222Q^ mice on Hi-B6 and WT mice on either diet.

### Pyridoxine supplementation blocks activation of stress signaling in SPL^R222Q^ mouse kidneys

Consistent with the effects on podocytes observed by STED microscopy, expression of podocyte markers nephrin and synaptopodin was reduced in extracts of SPL^R222Q^ kidneys after withdrawal of pyridoxine supplementation, as shown by IB (**Figure 6A-B**). To gain further insight into the molecular events leading to podocyte injury and nephrosis in the SPL^R222Q^ mice, we used IB of whole kidney tissue extracts to compare the four groups of mice for the activity of pathways known to be activated by S1P signaling and/or SPL insufficiency and implicated in the pathogenesis of fibrosis, including AKT (26, 27), ERK (27–29); PKCo (30), STAT3 (23), and SMAD3/4 (31). In each case, we observed upregulation of the phosphorylated (active) form of the signaling protein in the SPL^R222Q^ mouse kidneys upon pyridoxine withdrawal, but not in the three control groups. STAT3 and S1P signaling pathways are mutually co-activating (32–34). In mice and humans, STAT3 activation has been linked to diabetic kidney disease and other forms of glomerular injury (35–37). Hence, we examined the STAT3 pathway in greater detail. Performing p-STAT3 IHC, we observed increased signals of p-STAT3, especially in the tubule cells and interstitium of SPL^R222Q^ mouse kidneys after withdrawal of pyridoxine supplementation, but not in the control groups (**Figure 7A**). Moreover, qPCR confirmed that downstream targets of STAT3 transcriptional regulation, including *Lcn2, Timp1, Socs1,* and *Socs3*, were upregulated in SPL^R222Q^ mice kidney after withdrawal of pyridoxine supplementation (**Figure 7B**).

**Figure 6.**
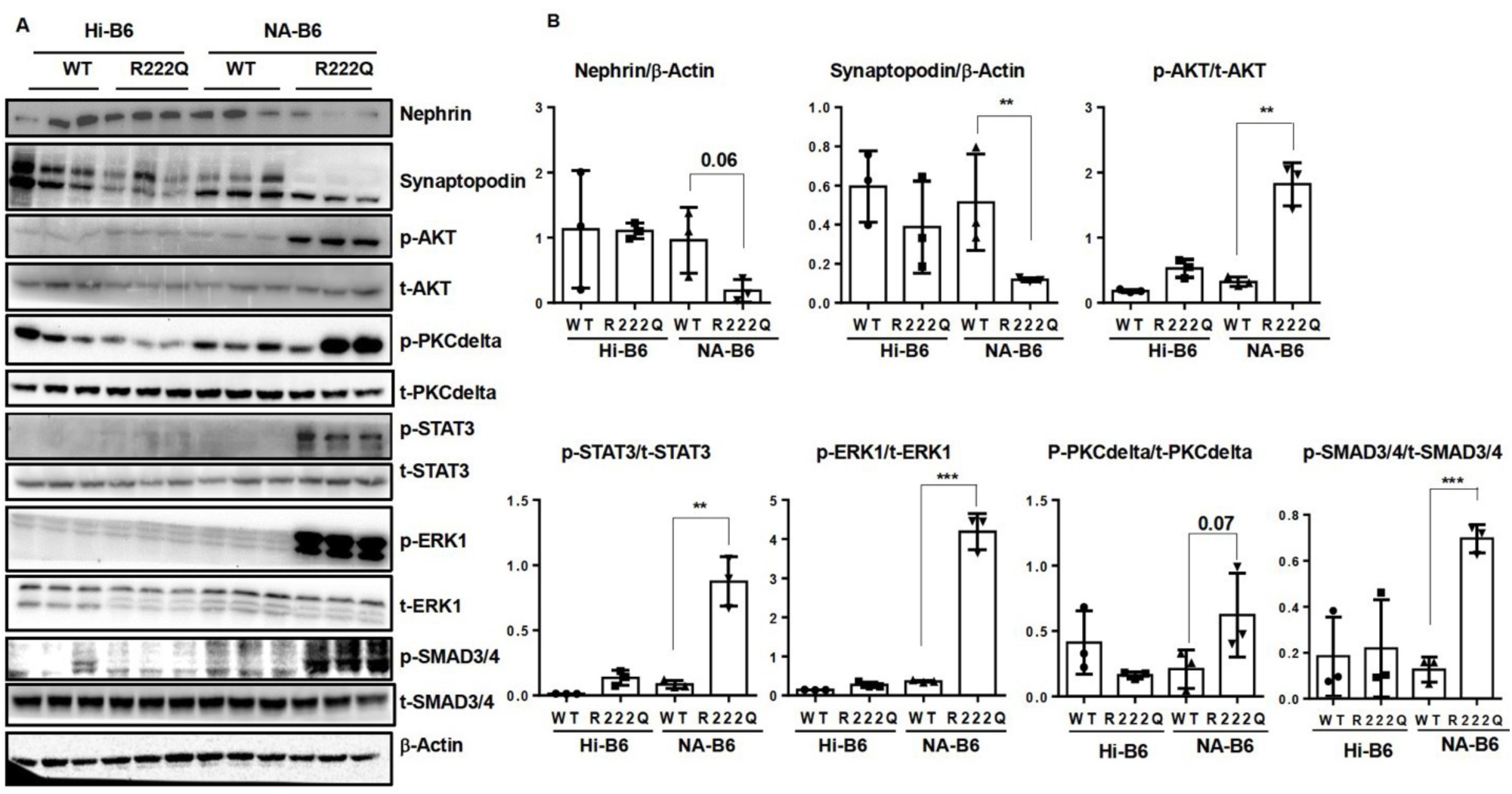
Pyridoxine supplementation prevents activation of stress signaling in SPL^R222Q^ mouse kidneys. (A) IB showing that podocyte markers nephrin and synaptopodin were reduced in the kidneys of SPL^R222Q^ mice on NA-B6 diet compared to SPL^R222Q^ mice on Hi-B6 and WT mice on either diet. Hyperactivation of p-AKT, p-PKC delta, p-STAT3, p-ERK1, and p-SMAD are observed in the kidney of SPL^R222Q^ mice on NA-B6 chow. D-actin is a loading control. (B) Quantification of corresponding IB. * p ≤ 0.05, ** p ≤ 0.01, and *** p ≤ 0.001.

**Figure 7.**
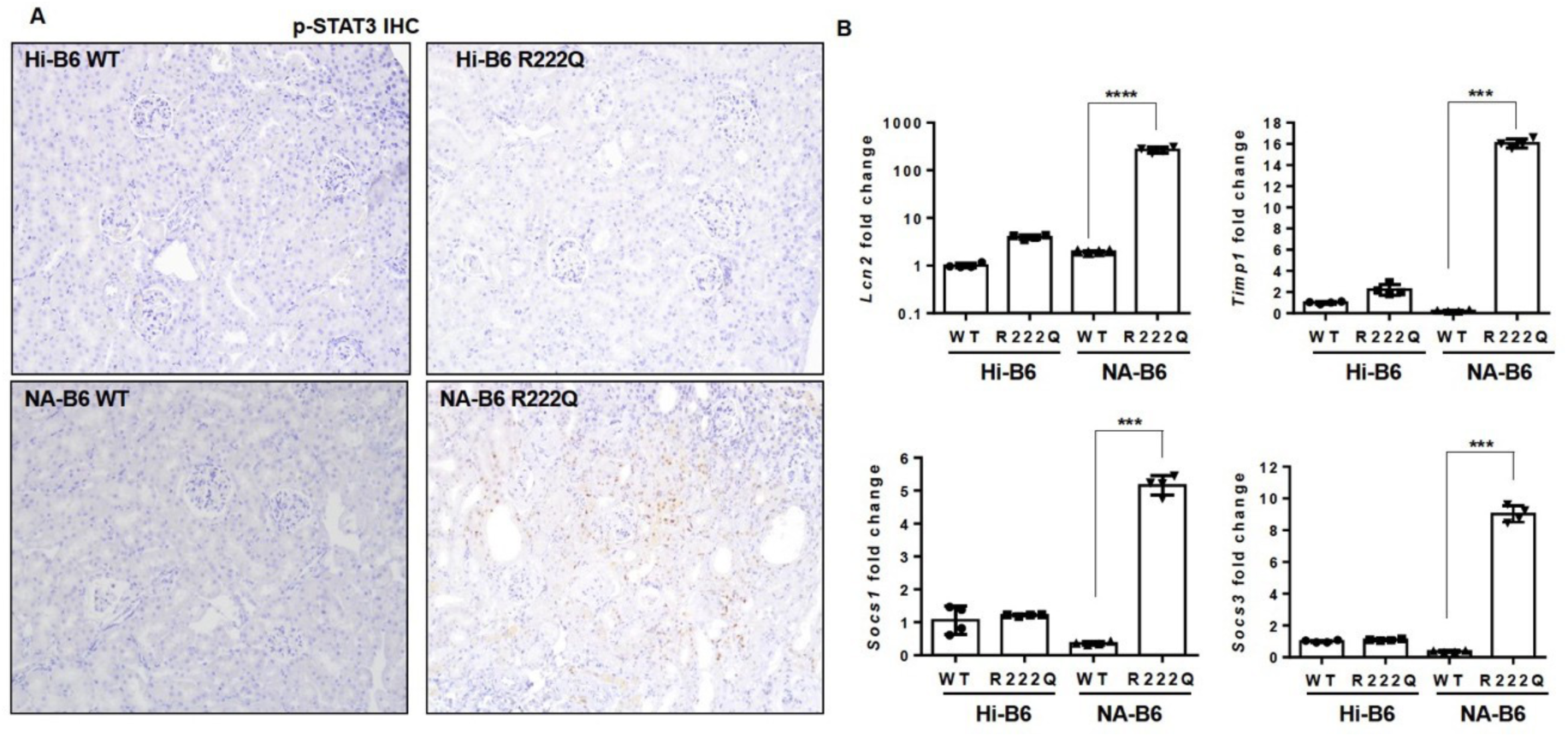
Low levels of pyridoxine cause activation of p-STAT3 and its target genes in the kidney of SPL^R222Q^ mice. (A) p-STAT3 IHC staining displaying more staining in kidney sections of SPL^R222Q^ mice fed NA-B6 chow compared to SPL^R222Q^ mice on Hi-B6 and WT mice on either diet. (B) Relative expression of STAT3 target genes *Lcn2, Timp1, Socs1*, and *Socs3* (normalized with D-actin) in SPL^R222Q^ mice fed NA-B6 chow compared to SPL^R222Q^ mice on Hi-B6 and WT mice on either diet (n = 4). * p ≤ 0.05, ** p ≤ 0.01, *** p ≤ 0.001, and **** p ≤ 0.0001.

### Pyridoxine supplementation suppresses a transcriptional profile reflecting tissue injury and extracellular matrix (ECM) remodeling in SPL^R222Q^ mouse kidneys

To assess the impact of SPL insufficiency on kidney homeostasis, we conducted a transcriptome analysis using RNA sequencing (RNA-Seq) of whole kidney tissues from WT and SPL^R222Q^ mice on Hi-B6 chow and three weeks after switching to NA-B6 chow. The Pearson correlation and principal component analysis revealed a distinct and significantly altered expression pattern of kidney transcripts in SPL^R222Q^ mice transferred to NA-B6 chow not observed in the other three groups, as shown in **Supplementary Figures S2**, and depicted using heat maps and overall transcriptome profile (**Supplementary Figure S3** and **S4**). To focus on differentially expressed genes (DEGs) whose transcription levels were specifically affected by SPL status, we excluded all genes that were differentially expressed in response to pyridoxine alone, i.e., between WT Hi-B6 versus WT NA-B6 groups. This led to the exclusion of nine genes (*Hdc* encoding histidine decarboxylase, the cytochrome p450 genes *Cyp7b1* and *Cyp4a14, Spp1* encoding osteopontin, the fibronectin gene *Fn1, Calb1* encoding calbindin 1, *Pigr* encoding the polymeric immunoglobulin receptor, *Chd9* encoding chromodomain helicase DNA binding protein 7, and *Slc7a13* encoding the cystine transporter). Of these, only histidine decarboxylase is known to be dependent upon pyridoxal 5’-phosphate for function. The remaining gene set is shown in **Figure 8A**. Of note was the fact that kidney injury molecule 1 (*Kim1/Havcr1*), an immunoglobulin superfamily protein and established indicator of kidney injury, was upregulated in SPL^R222Q^ mice on NA-B6 chow (38). GO enrichment analysis revealed many other related processes that were highly enriched by SPL insufficiency. Pathways involved in the innate immune response including chemokine-and cytokine-mediated signaling (*Osrm, Ccr5, Cx3Cr1, Cxcl15, C3ar1*), TNF-a signaling (*Gadd45g, Fam129a* and *Hsph1*), macrophage activation (*Cd68*), and complement activation (*Pzp, Hrg, Serpina16, Wfdc8, Spint4, Tmprss11a*) were differentially expressed (all upregulated with the exception of *Cxcl15*), consistent with their roles in the pathogenesis of kidney injury and kidney fibrosis (39–41). Upregulation of genes related to IL-17 (*Mmp2, Mmp14*), a known STAT3 transcriptional target, is consistent with our findings of STAT3 activation in kidneys of SPL^R222Q^ mice on NA-B6. (42). An interconnected signaling network that includes interactions between TGFD/SMAD, YAP/TAZ, IGF and HIPPO pathways central to tissue repair and fibrosis (*Igfbp1, Grem2, Stc1, St2, Pappa, Axl, Adamtsl2*) were differentially expressed (all upregulated with the exception of *Adamtsl2*) (43). Eleven genes representing collagen fibril organization and extracellular matrix (ECM) remodeling pathways directly involved in the development of fibrosis were dysregulated in SPL deficient kidneys (*Col12a1, Col1a1, Col1a2, Dcn, Ddr2, Adamtsl2, Lamc3, Zfp469, Dsg3, MMP2, MMP14*), all of which were upregulated, with exception of *Lamc3* and *adamtsl2*. This indicates that injury signals leading to maladaptive tissue repair were already present in the kidney after three weeks of low B6 conditions, well before pathological signs of fibrosis could be detected (31, 44). In addition, cholesterol metabolism (*Apoa4, Apob, Azgp1*) and lipid metabolic pathways that play roles in podocyte injury, kidney disease, and which interact closely with sphingolipids were upregulated (45–47). Protein degradation, which is known to be influenced by SPL through the endoplasmic reticulum stress response pathway and other mechanisms, was dysregulated in SPL deficient kidney (13, 48). Response to infections with *Staphylococcus aureus, Influenza A, Entamoeba histolytica* (which causes amoebiasis) and *Plasmodium* (the malaria parasite) were functionalities represented by genes corresponding with immune, ECM and cytokine pathways mentioned above. (**Figure 8B**). Six genes involved in vascular smooth muscle contraction (*Slc8a1, Gpr176, Scn2b, Clca3a1, Cacna1s, Pvalb*) were upregulated, which may be explained by the vasoconstrictive effect of S1P on pre-glomerular microvessels, especially afferent arterioles (49). Genes involved in arginine and proline metabolism are upregulated likely because of the known role of PLP in conversion of arginine to proline and, thus, may not be relevant to loss of SPL function independent of pyridoxine status. Transcriptomic profiling of mutant mouse kidneys identified the chemokine signaling pathway as one of the prominently activated pathways. Consistent with KEGG pathway analysis (**Supplementary Figure S5)**, several major downstream effectors of chemokine signaling, including STAT3, ERK1, AKT, and Ca²⁺-associated signaling (PKC), were implicated in the SPL^R222Q^ mice kidneys. These RNA-seq findings were consistent with our IB results, which demonstrated increased activation of STAT3, ERK1, and AKT, along with enhanced calcium-related signaling. Taken together, these results support the conclusion that chemokine signaling is activated in mutant kidneys and may play an important role in driving renal inflammatory and downstream cellular responses. Using qPCR to measure gene expression, we confirmed the upregulation of key transcripts, including established kidney injury markers (*Kim1/Havcr1, Pappa)*, chemokines and their receptors (*Cxcr1, Ccl17, Ccl2/Mcp1, Cxcl10*), and pro-inflammatory and profibrotic cytokines (*Tnfa* and *Tgfb*) (**Figure 8C**). In addition, we observed upregulation of *Tnfsf15*, a cytokine recently linked to non-monogenic steroid-sensitive nephrotic syndrome (50). Altogether, kidney transcriptional profiling confirms that a process of pro-inflammatory signaling and extracellular matrix remodeling, ultimately leading to glomerulosclerosis and podocyte injury, is induced within weeks after discontinuation of pyridoxine supplementation in SPL^R222Q^ mice.

**Figure 8.**
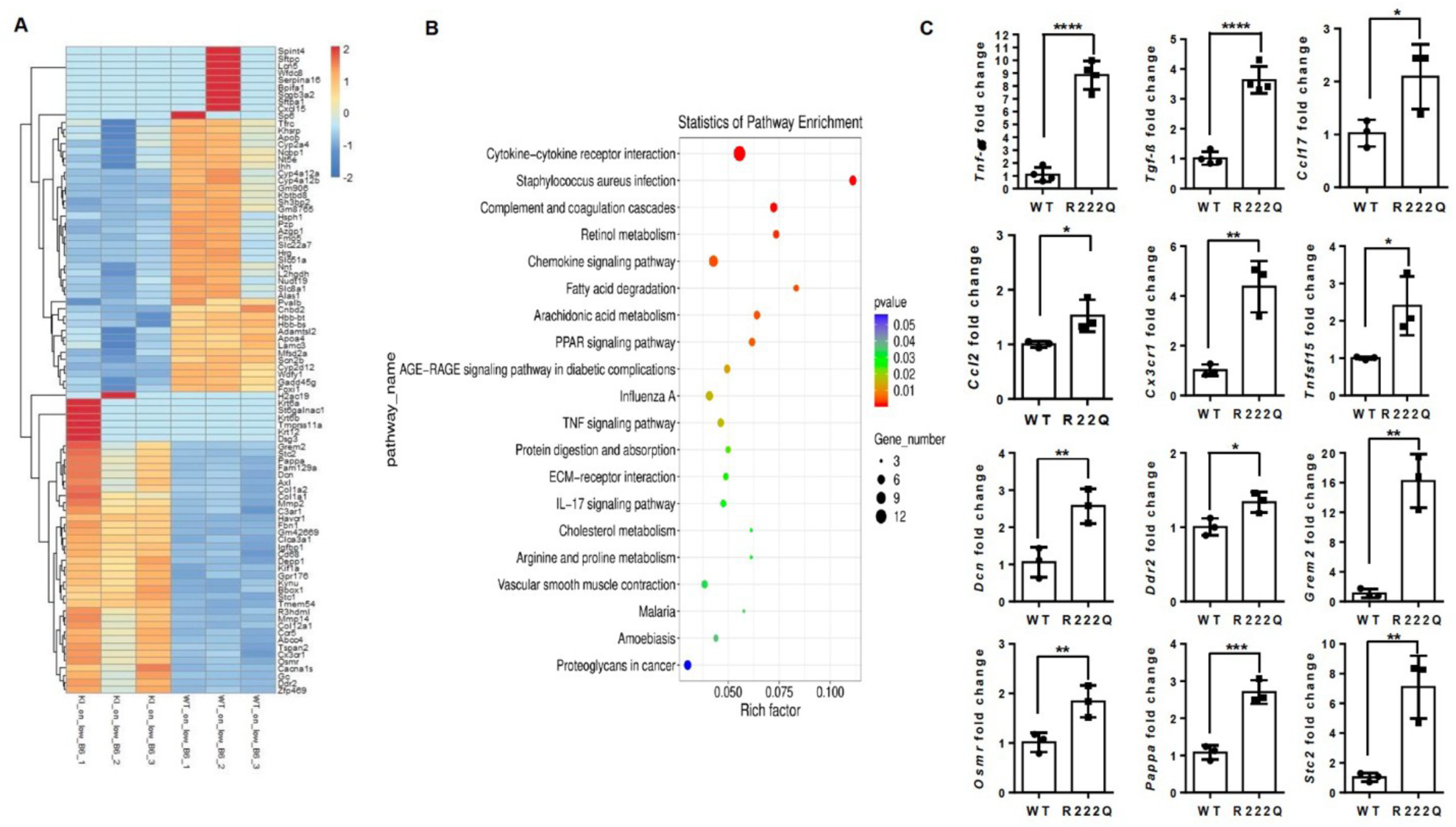
RNAseq analysis and qPCR validation of target genes. (A) Heatmap displaying upregulation of chemokine, cytokine, extracellular matrix remodeling, and collagen-related genes in SPL^R222Q^ mice versus WT mice fed NA-B6 chow. (B) Statistics of pathway enrichment further confirm the aberrant regulation of chemokine, cytokine, immune, TNF and other signaling pathways. (C) qPCR validation of target genes. At least one to two target genes were validated from each upregulated pathway. * p ≤ 0.05, ** p ≤ 0.01, and *** p ≤ 0.001.

### SPLIS nephrosis is caused by S1P accumulation

The development of SPLIS phenotypes in SPL^R222Q^ mice receiving limiting amounts of B6 in their diets coincided with a striking rise in circulating and tissue S1P levels, thus implicating S1P in the pathogenesis of the disease. This notion was further supported by the upregulation of S1P-linked signaling pathways in their kidneys. However, the pathogenic role of SPL product deficiencies and/or accumulation of other sphingolipids could not be ruled out. To test whether S1P accumulation mediates the SPLIS nephropathy, we placed SPL^R222Q^ mice on NA-B6 chow for two months, followed by two weeks of parenteral administration of SKI II, a broad-spectrum agent which has been shown to inhibit the catalytic activity of both isoforms of sphingosine kinase, SphK1 and SphK2 and to additionally induce proteolytic degradation of SphK1. Treatment of SPL^R222Q^ mice with SKI II resulted in the complete rescue of kidney barrier function and overall kidney health, as shown by normalization of urine ACR, serum albumin, BUN and creatinine to levels indistinguishable from WT (**Figure 9 A-D**). Treatment of SPL^R222Q^ mice with the selective SphK1 inhibitor PF543 under the same conditions gave nearly the same result, indicating that the pool of S1P responsible for inducing kidney injury was largely, if not completely, generated by SphK1 (**Figure 9 A-D**).

**Figure 9.**
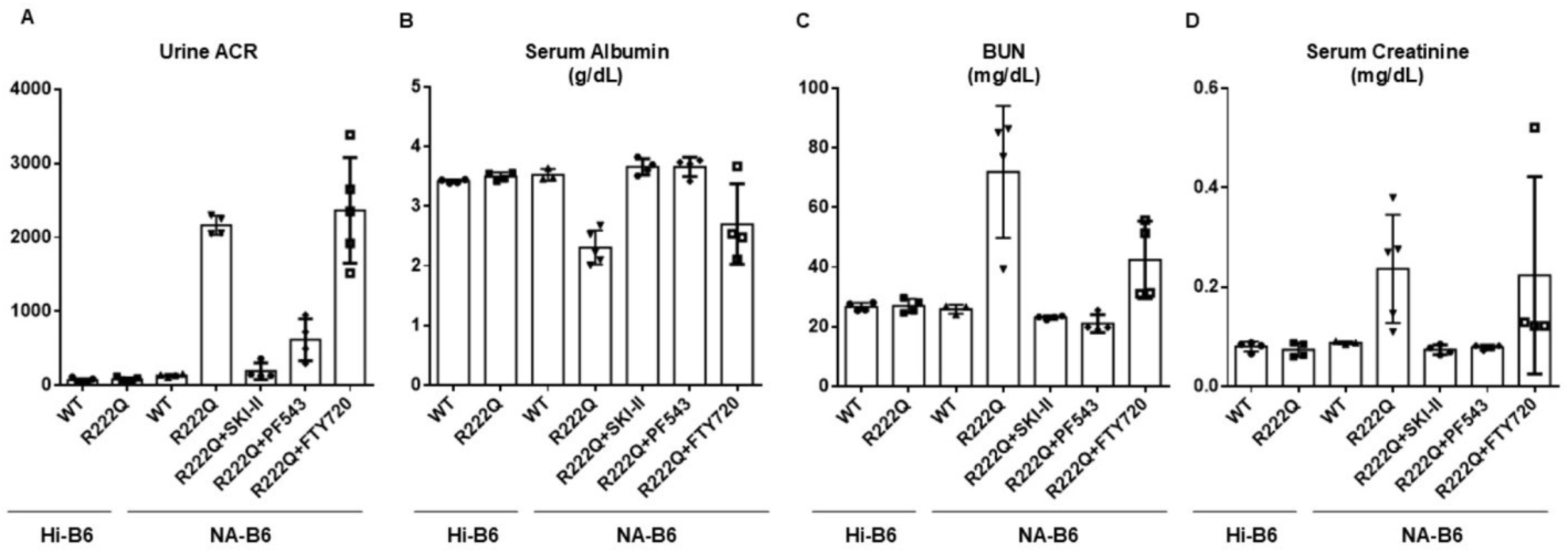
Inhibition of S1P production prevents SPLIS nephropathy. Parameters of kidney health were compared in WT and SPL^R222Q^ mice on Hi-B6 or NA-B6 diets versus SPL^R222Q^ mice on NA-B6 and simultaneously treated with either S1P receptor antagonist FTY720, nonspecific SphK inhibitor SKI II, or SphK1-specific inhibitor PF543. (A) Urine ACR, p < 0.0001. SPL^R222Q^ vs SPL^R222Q^ treated with FTY720, NSD. (B) Serum albumin. SPL^R222Q^ vs SPL^R222Q^ treated with SKI II and SPL^R222Q^ treated with PF543, p < 0.0001. SPL^R222Q^ vs SPL^R222Q^ treated with FTY720, p = 0.342. (C) Blood urea nitrogen (BUN). SPL^R222Q^ vs SPL^R222Q^ treated with SKI II, p = 0.004, and SPL^R222Q^ vs SPL^R222Q^ treated with PF543, p = 0.0039. SPL^R222Q^ vs SPL^R222Q^ treated with FTY720, NSD. (D) Serum creatinine. SPL^R222Q^ vs SPL^R222Q^ treated with SKI II, p = 0.02, and SPL^R222Q^ vs SPL^R222Q^ treated with PF543, p = 0.03. SPL^R222Q^ vs SPL^R222Q^ treated with FTY720, NSD.

To determine whether S1P is acting through S1P receptors to mediate SPLIS kidney injury, SPL^R222Q^ mice were given NA-B6 chow for two months and then treated for two weeks with parenteral FTY720, which functionally antagonizes S1P1, S1P3, S1P4 and S1P5. In contrast to the kidney protective effect afforded by sphingosine kinase inhibition, blocking S1P signaling with FTY720 did not prevent the development of SPLIS nephrosis in SPL^R222Q^ mice (**Figure 9 A-D**). Taken together, these results support the conclusion that a SphK1–S1P pathway likely mediated by intracellular S1P functions of S1P drives kidney injury in SPLIS.

## Discussion

SPLIS is a recently recognized inborn error of metabolism predicted to afflict thousands of individuals worldwide. Like many monogenic forms of steroid-resistant nephrotic syndrome, SPLIS-associated nephrosis usually progresses rapidly to renal failure and is, directly or indirectly, the primary cause of childhood deaths in SPLIS (4). While kidney transplantation can have therapeutic benefit in SPLIS patients with end-stage kidney disease, the procedure is invasive, requires long-term immunosuppression, and many SPLIS children are not suitable surgical candidates (4). Further, kidney transplantation may not address the extrarenal manifestations of SPLIS. Alternative therapies are direly needed, particularly targeted treatments that address the root cause of SPLIS and have the potential to prevent or delay kidney failure and, ideally, other manifestations of this heterogeneous condition.

Vitamin B6 supplementation has been used successfully to treat deficiencies of enzymes that rely on PLP as a cofactor. Due to its phosphorylated state, PLP cannot be taken up readily by cells. However, cells can take up the unphosphorylated B6 vitamer pyridoxine and convert it intracellularly to PLP. Supplementation with pyridoxine increases intracellular PLP levels, thereby overcoming the reduced cofactor binding affinity and/or improving the protein stability of some pathogenic PLP-enzyme variants. Pyridoxine is given orally and exhibits an acceptable safety profile in infants and pregnant women (51). Despite these attributes, pyridoxine’s therapeutic activity is limited to a subset of variants in each inborn error of metabolism. This situation necessitates testing variants (or patients) individually. Further, pyridoxine treatment at doses greater than 500 mg/day may result in a sensory peripheral neuropathy (52). Demonstration that pyridoxine supplementation affords metabolic correction and disease-modifying activity in preclinical SPLIS models would provide the rationale for long-term pyridoxine treatment and monitoring.

In this study, we employed three complementary approaches to establish proof-of-concept for the efficacy of cofactor supplementation in SPLIS. We focused on the SPL R222Q variant, as it is the most common SPLIS-associated pathogenic variant, and its low expression, activity, and predicted cofactor binding affinity make it a candidate for both the prosthetic and chaperone functions of PLP (4, 20). Further, we had previously observed clinical improvement in a SPLIS patient homozygous for the SPL R222Q variant in response to pyridoxine. Additionally, we have observed the restoration of sphingolipid metabolism in R222Q patient-derived skin fibroblasts (20). Here, we show that the peripheral neuropathy of a second patient with R222Q variant SPLIS abated after pyridoxine administration, objectively measured by NCS. This occurred concomitant with a reduction in the patient’s plasma S1P levels and was consistent with a significant rise in SPL activity after pyridoxine treatment in patient-derived skin fibroblasts. Furthermore, the purified recombinant human SPL R222Q variant exhibited a dose-dependent increase in catalytic activity in response to increasing concentration of PLP. In contrast, PLP exerted no effect on the R222W variant, in which the aromatic tryptophan substitution is predicted to cause a more severe structural distortion than the glutamine substitution at the same residue (53). Additional PLP also had no effect on the WT enzyme, which binds PLP optimally during polypeptide synthesis and assumes a thermodynamically favorable (native) protein conformation. Thus, the SPL holoenzyme would not be expected to benefit catalytically or be further stabilized by cofactor supplementation. Thus, our *in vitro* findings demonstrate the R222Q variant requires higher concentration of PLP for its binding during catalytic activity than the WT enzyme does. Although pyridoxine only raises SPL R222Q-variant enzyme activity to about 25% of healthy control levels, this amount of activity has been shown to prevent the consequences of SPL insufficiency in animal models (21). This result establishes the biochemical basis for pyridoxine supplementation in SPL R222Q-variant SPLIS.

To further explore the impact of pyridoxine administration on tissue SPL activity, sphingolipid metabolism, organ function, and pathology in SPLIS, we used gene editing to generate the SPL^R222Q^ mouse, a customized knock-in animal model of SPL R222Q-variant SPLIS. Despite the low SPL activity observed in tissues of SPL^R222Q^ mice fed Hi-B6 chow, which ranged from 25-50% of WT levels, they did not accumulate S1P, indicating this level of enzyme activity is sufficient for proper sphingolipid metabolism. Similarly, none of the SPLIS-associated phenotypes were detected under these conditions. However, upon withdrawal of pyridoxine supplementation, SPL^R222Q^ mice (but not WT mice) showed metabolic, neurological, renal and hematological deterioration. Thus, the SPL^R222Q^ mouse represents a novel and inducible SPLIS model with many advantages over the *Sgpl1* KO model. Because of its ability to reach adulthood, the conduct of complex procedures and assessment of the efficacy of therapeutic agents that act by modifying the activity or stability of the variant are possible in SPL^R222Q^ mice. Our results simultaneously confirmed that the SPL-R222Q variant is pyridoxine responsive, and that progression of kidney disease should be preventable in SPLIS patients harboring pyridoxine-responsive pathogenic variants of SPL.

Histological and molecular methods combined with transcriptional profiling were used to characterize the impact of SPL insufficiency on murine kidney. Our kidney transcriptome results demonstrate that inactivation of the SPL-R222Q variant led rapidly to kidney injury, as evidenced by upregulation of *Kim1/Havcr1*, a well-established marker of kidney injury which is also reported to influence lipid uptake in proximal tubule cells (38). Loss of SPL function also led to activation of the innate immune system including effects on TNFa, IL-17, TGFD/SMAD, and other chemokines and cytokines, macrophage polarization, and complement activation. A prominent signature of ECM remodeling presaged the later development of kidney fibrosis. Our immunoblotting results showed that SPL insufficiency led to activation of STAT3, AKT, MAPK/ERK, PKCo and TGFD, stress pathways that are known downstream effectors of both cytokines and S1P/SPL. Pathological results confirmed the essential role of SPL in kidney homeostasis as demonstrated by podocyte injury, podocyte loss, glomerular injury, protein casts, and kidney fibrosis when SPL was inactivated.

S1P is generated through phosphorylation of the precursor sphingosine by either SphK1 or SphK2. SphK1 is largely cytosolic and can translocate to the plasma membrane upon stimulation, producing a pool of S1P that can be exported to engage S1P receptors in an autocrine/paracrine manner (54). SphK2 is enriched in intracellular compartments (nucleus, endoplasmic reticulum and mitochondria) and is frequently linked to regulation of intracellular S1P pools that can influence transcriptional programs and stress responses (54). By inhibiting S1P production using either a nonspecific or selective inhibitor of SphK1, kidney protection was afforded to SPL^R222Q^ mice under conditions that uniformly lead to SPLIS nephropathy. We observed a modest increase in efficacy of the nonspecific inhibitor over the selective SphK1 inhibitor PF543 based on ACR normalization. There are several possible explanations for this. First, the nonselective inhibitor SKI II also impacts the activities of SphK2 and dihydroceramide desaturase (55). These enzymes could potentially contribute to SPLIS nephropathy in a minor way. A more likely explanation is the fact that SKI II induces proteolytic degradation of SphK1, which could have a more lasting and profound effect on SphK1 inhibition beyond its catalytic effects (56). Regardless of these possibilities, the major effect is clearly mediated by an SphK1-generated pool of S1P. S1P mediates many of its biological effects through activation of its receptors (57). However, the broad spectrum S1P receptor antagonist FTY720 did not prevent SPLIS nephropathy. This could be due to S1P’s receptor-independent functions, such as calcium signaling, epigenetic regulation of gene expression through its effect on histone deacetylases, DNA repair and telomere maintenance, and mitochondrial function (58–60). Alternatively, the effect could be mediated through activation of S1P_2_, the only S1P receptor not antagonized by FTY720. Future studies will be required to determine this.

Some caveats to our study should be noted. We did not carry out the study for an extended period. Thus, we cannot rule out the possibility that kidney disease or other phenotypes might eventually develop in SPL^R222Q^ mice under Hi-B6 conditions. Some SPLIS phenotypes, such as glucocorticoid deficiency, hypercholesterolemia, and reproductive health, were not assessed in our model. These will be evaluated in future studies. Clinical studies testing the efficacy of pyridoxine will be needed to validate our preclinical findings. Importantly, although SPL R222Q represents the most prevalent pathogenic SPL variant in SPLIS, many other variants have been identified, including many missense variants with the potential to be activated by pyridoxine supplementation (4). Experimental testing of these variants will be necessary before extrapolating our findings. Our unpublished results suggest that at least four other SPLIS-associated SPL variants are responsive to pyridoxine, the detailed findings of which are out of the scope of this study.

In conclusion, our findings using a personalized murine knock-in mouse model of SPL^R222Q^ SPLIS demonstrate that pyridoxine supplementation improves tissue SPL activity, reduces tissue and plasma S1P levels to near normal levels, and prevents nephrosis, glomerulosclerosis, and other SPLIS phenotypes.

## Materials and Methods

### Sex as a variable

In this study, plasma S1P levels in one female subject with the rare disease SPLIS were compared to the S1P levels in healthy controls of both sexes. Our unpublished findings indicate that plasma S1P levels in subjects with SPLIS regardless of sex are at least two standard deviations above the mean compared to healthy subjects of both sexes, and SPLIS incidence is equal in both sexes. In animal studies, we used male and female mice with roughly equal distribution in all treatment groups. Based on years of observation, we have no evidence that sex influences the development or severity of kidney, neurological or immunological phenotypes in SPLIS or SPLIS mouse models. Thus, we believe our findings are equally relevant to both sexes.

### WES

Blood samples, clinical data, and fibroblasts were obtained from the proband after obtaining informed consent. DNA for WES and plasma for sphingolipid analysis pre-and post-pyridoxine treatment were obtained from blood. The quality of the DNA was evaluated by 1% agarose gel electrophoresis. The DNA libraries for sequencing were constructed according to the manufacturer’s protocol. Subsequently, we conducted the sequencing on an Illumina HiSeq 2000 platform. Sequencing reads were aligned to the human genome (GRCh37/hg19). SAM files from each sample were then converted to BAM format, sorted, and merged using SAMtools (http://samtools.sourceforge.net/). PCR duplicates were removed from the files using Picard (http://picard.sourceforge.net/). Further processing was carried out utilizing the Genome Analysis Toolkit (GATK) from the Broad Institute (http://www.broadinstitute.org/gatk/). All BAM files underwent local realignment with an indel realigner. Single-nucleotide variants (SNVs) and insertions/deletions (indels) within the captured coding exonic regions were identified using GATK’s Unified Genotype. The detected variants were filtered according to the criteria that we previously reported (1). Only the *SGPL1* variant (c.887C>A, R222Q) was segregated in the proband, following a recessive inheritance pattern. The conservation of mutations was assessed using CLUSTAL W.

### Isolation and propagation of skin fibroblasts

Fibroblasts at the primary stage were isolated as described in (61) and transduced with *hTERT* through the lentivirus vector pLV-hTERT-IRES-hygro (Addgene plasmid # 85140). Hygromycin at a concentration of 100 μg/mL was introduced for selection after two days post-transduction. In some cases, retroviral vector pBABE-puro-hTERT (Addgene plasmid # 1771) and puromycin selection (at 1-2 µg/ml) were employed to establish immortalized fibroblasts. Following the identification of stable cells, the immortalized fibroblast lines were expanded and sub-cultured for subsequent experiments.

### SPL activity and sphingolipid quantification

Recombinant SPL activity was measured by reacting 10 µM S1P (Avanti Polar Lipids) in PBS with 50 nM purified enzyme and varying concentrations of PLP. Reactions were quenched by mixing 1:1 with 2X DAB buffer (2X: 500mM Sodium Bicarbonate pH 10.0, 2% SDS, 2.5mM β-mercaptoethanol, 2.5mM 1,2 diacetylbenzene (DAB, Sigma-Alrich) and fluorescence of DAB upon reaction with the SPL product ethanolamine phosphate was measured at excitation/emission wavelengths of 355nm/455nm after 1 hour. SPL activity measured in human skin fibroblasts and tissues of SPL^R222Q^ and mice by monitoring formation of the (2*E*)-hexadecenal product using liquid chromatography-tandem mass spectrometry (62). For quantification of S1P in murine tissues and plasma, lipid extraction was performed, followed by separation and identification using tandem mass spectrometry on an Agilent 6495C triple quadrupole instrument as described (63).

### NCS

NCS were conducted by utilizing a Dantec Keypoint Portable Nerve Conduction/Electromyography machine from Dantec Dynamics, Bristol, UK. The findings were analyzed and reported by a neurophysiologist, and conduction defects were categorized in accordance with the Neurophysiological Classification of Carpal Tunnel Syndrome.

### Construction of R222Q and R222W variants of the *SGPL1* gene

Recombinant WT *SGPL1* sequence was purchased as an *E. coli* codon-optimized gBlock from Integrated DNA Technologies. End primers were designed to remove the first 92 N-terminal residues and insert a C-terminal 6xHis tag with a GSG linker (**Table S2**). The insert was digested with NdeI and XhoI and ligated into the respective sites in the vector pET22b. R222 mutations (Q and W) were introduced using overlap extension PCR, and all constructs were confirmed by sequencing with T7 primers.

### Protein expression and purification

Wild-type, R222Q, and R222W SPL variants were introduced into *E. coli* BL21 (DE3) containing helper plasmid pG-KJE8 (Takara) and grown at 37°C before inducing with 0.1mM IPTG and 0.2% Arabinose and expressing overnight at 25°C. Expression cultures were pelleted and stored at-20°C until day of purification. Purification was performed as described previously (64). Briefly, cells were lysed in buffer containing excess PLP (20mM Tris, 300mM NaCl, 10mM Imidazole, 15% sucrose, 1mM PLP, 0.125mM PMSF, pH to 8.0 with HCl) and loaded on a Ni-NTA affinity column. Enzyme was eluted under low-salt (10mM NaCl) conditions and subsequently cleaned up by cation exchange chromatography (UNOsphereS, Bio-Rad). PLP was included in the initial step of purification to ensure formation of the holoenzyme but was excluded from later steps to determine the effect of supplementation *in vitro*.

### Animals

All animal studies were conducted under approved IACUC protocols at the NIH and at UCSF. To generate SPL**^R222Q^** mice, we introduced the R222Q mutation into exon 7 of the mouse Sgpl1 gene using CRISPR/Cas9 editing technology as described previously (65). We designed a single-guide RNA (sgRNA) containing a 20 bp target sequence, 5’ GATGGCCTGCAAAGCTTACC 3’ (Synthego), on mouse exon 7. To introduce the mutant sequence, we designed a single-stranded 199-base oligodeoxynucleotide (SPLIS ssOligo) containing the *SGPL1*c.665G>A SPLIS variant, flanked by 92 bases on the 5’side and 106 bases on the 3’ side: 5’ AGGCTGTACCTCCTGATACCCATCCTTAACTTACTCTGGTTTCTTTTCTTCACATAGG TGACTTCTGGGGGAACGGAAAGCATCCTGATGGCATGCAAAGCTTACCAGGACTTG GCGTTAGAGAAGGGGATCAAAACTCCAGAAATGTATGGATGTGTGTGTTTGTTTCCC TTCTGATATTGTCTATTTGTGGCAGCAC 3’ (IDT). The SPLIS ssOligo also contained a silent change to create a *SphI* restriction enzyme site for screening (**Figure 2**). The C57BL/6J strain was used as an embryo donor. Fertilized oocytes were injected with Cas9 protein (TrueCut Cas9 protein V2, Thermofisher), sgRNA and SPLIS ssOligo and transferred to the oviducts of pseudopregnant female mice. Founder mouse was confirmed by Sanger sequencing (**Figure 2A**). Mice were genotyped by PCR of tail-snip DNA at weaning using the flanking primer 1, 5’ AAGACTAGAATTGTAGG 3’ and primer 2, 5’ CCTCCTACAATTCTAGTCTT 3’ under the following PCR conditions: denaturation 95°C for 30 seconds, amplification 55°C for 1 minute, and extension 72°C for 1 minute (40 cycles). The resulting PCR product (443 bp) was digested with *SphI* to identify potential founders carrying the R222Q change. WT allele yielded a 443 bp, undigested fragment and the SPLIS ssOligo-modified allele yielded a 275 bp and 167 bp bands (**Figure 2B**). Mice were fed chow with Hi-B6 (18 mg/Kg) content or NA-B6 (AIN-93 chow with ∼2 ppm pyridoxine, LabDiet, San Francisco, CA, USA). Pyridoxine content was determined by the Nestle Purina Analytical Lab (St. Louis, MO, USA).

### IB

Kidneys from SPL^R222Q^ and WT mice on NA-B6 and Hi B6 chow were homogenized and cleared by centrifugation. The following antibodies for IB analysis were purchased from Cell Signaling Technology (CST, Danvers, MA, USA): anti-phosphorylated Stat3 (9145), anti–total STAT3 (9131), p-ERK1 (4370), t-ERK1 (4696), p-PKC delta (9376), t-PKC delta (2058), p-AKT (4060), t-AKT (9272), p-SMAD3 (9520) and t-SMAD3 (9513). Synaptopodin (SC-515842) and Nephrin (sc-376522) antibodies were purchased from Santa Cruz Biotechnology, CA, USA, while antibodies against murine SPL and β-actin were from Sigma-Aldrich (St. Louis, MO, USA). Antibodies against human SPL were from R&D Systems (Minneapolis, MI, USA). HRP-conjugated secondary antibodies [115-035-003, Goat Anti-Mouse IgG (H + L); 111-035-144, Goat Anti-Rabbit IgG (H + L), Jackson Immuno Research; sc-2020, donkey anti-goat IgG-HRP, Santa Cruz Biotechnology]. Signals were detected using the SuperSignal West Pico kit (Thermo Fisher Scientific). Radiographic bands were quantified using NIH ImageJ.

### Hematological and urine ACR analysis

Serum albumin and whole blood red blood indices were measured at University of California Davis Comparative Pathology Laboratory. Briefly, a total of 100 μL of diluted blood samples were analyzed using the XT-2000iV veterinary hematology analyzer (Sysmex, Kobe, Japan) to obtain a full standard hematology profile, while a COBAS INTEGRA 400 plus instrument (Roche Diagnostics, Indianapolis, IN, USA) was used to assess serum albumin levels.

### PAS staining and IHC

Kidney tissue samples were fixed in 4% paraformaldehyde with subsequent paraffin embedding and were sectioned to 5 µm thick. PAS and IHC were performed as we described previously (23, 66). Qualitative evaluation was performed by a nephropathologist to identify segmentally sclerosed glomeruli, glomeruli displaying mesangial hypercellularity, and interstitial fibrosis. For IHC, phospho-STAT3 (Tyr705) (D3A7) XP® rabbit monoclonal antibody (Cell Signaling, 9145, Danvers, MA, USA) was used. All images were captured using a Nikon Eclipse 80i microscope equipped with a digital camera (Nikon DS-Ri1, Tokyo, Japan).

### STED super-resolution fluorescent microscopy

STED was performed at the Optical Imaging Facility of USC Stem Cell on paraffin-embedded fixed kidney cortex sections. For antigen retrieval, heat-induced epitope retrieval with sodium citrate buffer (pH 6.0) was applied. To reduce nonspecific binding, sections were blocked with goat serum (1:20, Sigma-Aldrich, Cat# G9023). For tissue permeabilization, samples were incubated with 0.1% Triton X-100 for 10 min. Further samples were incubated for a multicolor staining with primary anti-nephrin (1:50, Progen, Cat#: GP-N2) and STED-compatible secondary Alexa Fluor 488 (1:500, Invitrogen, Cat# A-11073) antibodies. Samples were mounted in Prolong Diamond media (Invitrogen, Cat#: P36961) and covered with #1.5 glass coverslips. STED xyz images were acquired by using a Stellaris 8 STED microscope (Leica Microsystems) equipped with an HC PL APO 100x/1.4 OIL STEDWHITE objective. Pixel sizes of 20 to 25 nm were used for STED nanoscopy with line accumulation set to 4. The pinhole was set to 1.0 AU. Anti-nephrin-AF488 was excited at 492 nm wavelength, and STED was performed using a pulsed depletion laser at 592 nm wavelength. For both colors, acquisition time gating was set to 1 to 10 ns.

### TEM

Electron microscopic sample handling and detection were performed by the Core Center of Excellence in Nano Imaging (CNI) at the University of Southern California, Los Angeles, CA. The foot process width and the number of foot processes per μm of glomerular basement membrane were calculated using a curvimeter (Sakura Co., LTD, Tokyo, Japan). Five glomeruli were randomly selected from each mouse, and ten electron micrographs were taken and analyzed for each glomerulus.

### Grip strength testing

Each mouse underwent three sets of grip strength tests using a Chatillon Ametek digital force gauge model DFIS-10, as described previously (23).

### RNA extraction and RNA-Seq

Total RNA was extracted from kidney tissues using TRIzol reagent (Thermo Fisher Scientific, Waltham, MA, USA) according to the manufacturer’s protocol. RNA integrity was verified using an Agilent Technologies 2100 Bioanalyzer. Some RNA was used for qPCR described below. For RNA-Seq studies, ribosomal RNA was removed, followed by fragmentation using divalent cation buffers at elevated temperatures. The sequencing library was prepared following Illumina’s TruSeq-stranded-total-RNA-sample preparation protocol. Quality control analysis and quantification of the sequencing library were performed using Agilent Technologies 2100 Bioanalyzer High Sensitivity DNA Chip. Paired-end sequencing was performed on Illumina’s NovaSeq 6000 sequencing system.

### Transcript assembly and differential expression analysis of transcripts

RNA-Seq was commercially performed by LC Sciences, Texas, USA (https://lcsciences.com/). Cutadapt (https://cutadapt.readthedocs.io/en/stable/) and customized Perl scripts were used to remove the reads that contained adaptor contamination, low-quality bases, and undetermined bases. The sequence quality was verified using FastQC (http://www.bioinformatics.babraham.ac.uk/projects/fastqc/). We used HISAT2 (67) to map reads to the genome of *Mus musculus* (Version: v101). The mapped reads of each sample were assembled using StringTie (68). Then, all transcriptomes from 12 samples were merged to reconstruct a comprehensive transcriptome using Perl scripts and gffcompare (https://github.com/gpertea/gffcompare/). After the final transcriptome was generated, StringTie (68) and ballgown (http://www.bioconductor.org/packages/release/bioc/html/ballgown.html) were used to estimate the expression levels of all transcripts. StringTie was used to calculate the expression level of mRNAs and lncRNAs by calculating FPKM. mRNAs/lncRNAs differential expression analysis was performed by DESeq2 software (69) between two different groups (and by edgeR between two samples) (70). The genes/mRNAs/lncRNAs with FDR below 0.05 and absolute fold change ≥ 2 were considered differentially expressed genes/mRNAs/lncRNAs.

### GSEA

We performed gene set enrichment analysis using GSEA software (v4.1.0) (71) and MSigDB to identify whether a set of genes in specific GO terms, KEGG pathways, shows significant differences between the two groups. Enrichment scores and p-value were calculated using the default parameters. GO terms, KEGG pathways meeting this condition with |NES|>1, normalized enrichment score, p-val<0.05, FDR q-val<0.25 were considered different in the two groups. Gene Ontology (GO) (http://www.geneontology.org) terms were calculated using the Hypergeometric equation. GO terms with p-value <0.05 were defined as significant.

### qPCR

cDNA synthesis was performed by using the SuperScript III reverse transcriptase (Thermo Fisher Scientific). qPCR was performed using PowerUp SYBR Green Master Mix in QuantStudio™6 7 Flex Real-Time PCR System (Life Technologies, Thermo Fisher Scientific). Cq values of samples were normalized to the corresponding Cq values of *Actb*. The comparative Cq method was used to determine the quantification of the fold change in gene expression. Primers are listed in **Table S2**.

### S1P receptor blocking and sphingosine kinase inhibitors

SPL^R222Q^ mice were given NA-B6 chow for two months. Additional groups of WT mice given Hi-B6 and NA-B6 and SPL^R222Q^ mice given Hi-B6 for two months were used as controls. SPL^R222Q^ mice on NA-B6 were then treated for two weeks with either the S1P_1,3,4,5_ receptor antagonist FTY720 (Cayman Chemical) at 0.5 mg/kg, the specific SphK1 inhibitor, PF543 (Selleck Chemicals, LLC) at 1 mg/kg, the broad spectrum SphK1/2 and dihydroceramide desaturase inhibitor SKI II at 50 mg/kg (Cayman Chemical) or vehicle (DMSO in normal saline) given i.p. every other day. All agents were dissolved in DMSO stock solution. After the last dose, a 24h urine collection was obtained. Two days after the last dose, the mice were euthanized with terminal phlebotomy for serum albumin, creatinine and blood urea nitrogen (BUN). Both sexes were used for all treatment groups.

### Statistics

Data analysis was conducted with GraphPad Prism 5 (GraphPad Software). Two-tailed Student’s t-tests were employed to compare two groups, while ANOVA along with Bonferroni’s corrections were applied to compare multiple groups. A p-value of less than 0.05 was considered statistically significant.

### Study approval

Written informed consent was received prior to human subject participation in the study. Informed consent was obtained in accordance with an approved UCSF IRB protocol. Healthy pediatric and young adult controls undergoing elective surgeries for benign conditions were recruited from patients in the outpatient surgery center of UCSF Benioff Children’s Hospital, Oakland. Any patients with malignant, infectious, metabolic, genetic, hemolytic, autoimmune, or endocrine disorders were excluded. Plasma samples with visible hemolysis were also excluded. All animal protocols were approved by the Institutional Animal Care and Use Committee at UCSF and USC.

## Data availability

All RNA-Seq data for the article can be accessed in the Supplementary Files and the Gene Expression Omnibus repository.

## Author contributions

JDS designed the research studies, analyzed the data and, together with RK, wrote the manuscript. RK, MLA, EK, JYL, MRS, AI, VB, GG, JPP, RAK, GT, BO, RLP, and RZ, conducted experiments, acquired and analyzed data, provided reagents and edited the manuscript. All authors approved the final version of the manuscript.

## Abbreviations

ACR: Albumin creatinine ratio
BUN: blood urea nitrogen
DAB: 1,2 diacetylbenzene
ECM: extracellular matrix
GO: gene ontology
GATK: genome analysis toolkit
gRNA: guide RNA
Hi-B6: high B6
NCS: nerve conduction study
NA-B6: no added B6
PAS: periodic acid-Schiff
PLP: pyridoxal 5’-phosphate
RNA-Seq: RNA sequencing
SNVs: single nucleotide variants
ssODNs: single strand oligodeoxynucleotides
S1P: sphingosine-1-phosphate
SPL: sphingosine phosphate lyase
SPLIS: sphingosine phosphate lyase insufficiency syndrome
SRNS: steroid-resistant nephrotic syndrome
STED: stimulated emission depletion
TEM: transmission electron microscopy
WES: whole exome sequencing

---

**Table S1.** Quantification of glomerulosclerosis.

| **Genotype** | **Total**<br>**Glomeruli** | **Focal segmental**<br>**glomerulosclerosis**<br>**(%)** | **Global**<br>**glomerulosclerosis**<br>**(%)** |
| --- | --- | --- | --- |
| WT Hi-B6 (n = 3) | 241 | 0 | 0 |
| WT NA-B6 (n=3) | 175 | 0 | 0 |
| SPL<sup>R222Q</sup> Hi-B6 (n=3) | 230 | 0 | 0 |
| SPL<sup>R222Q</sup> NA-B6 (n=3) | 299 | 14.2 | 8.56 |

**Table S2.**
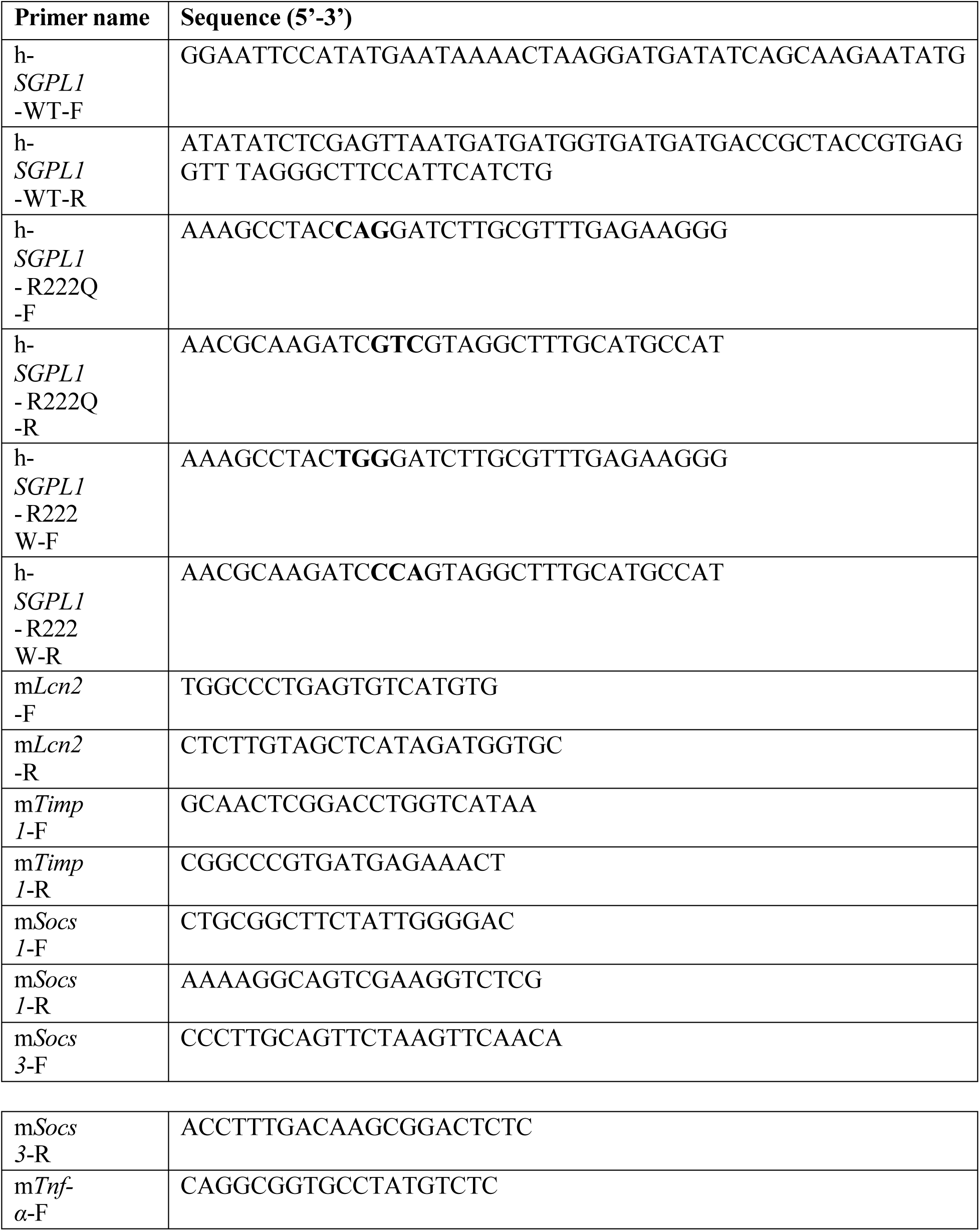
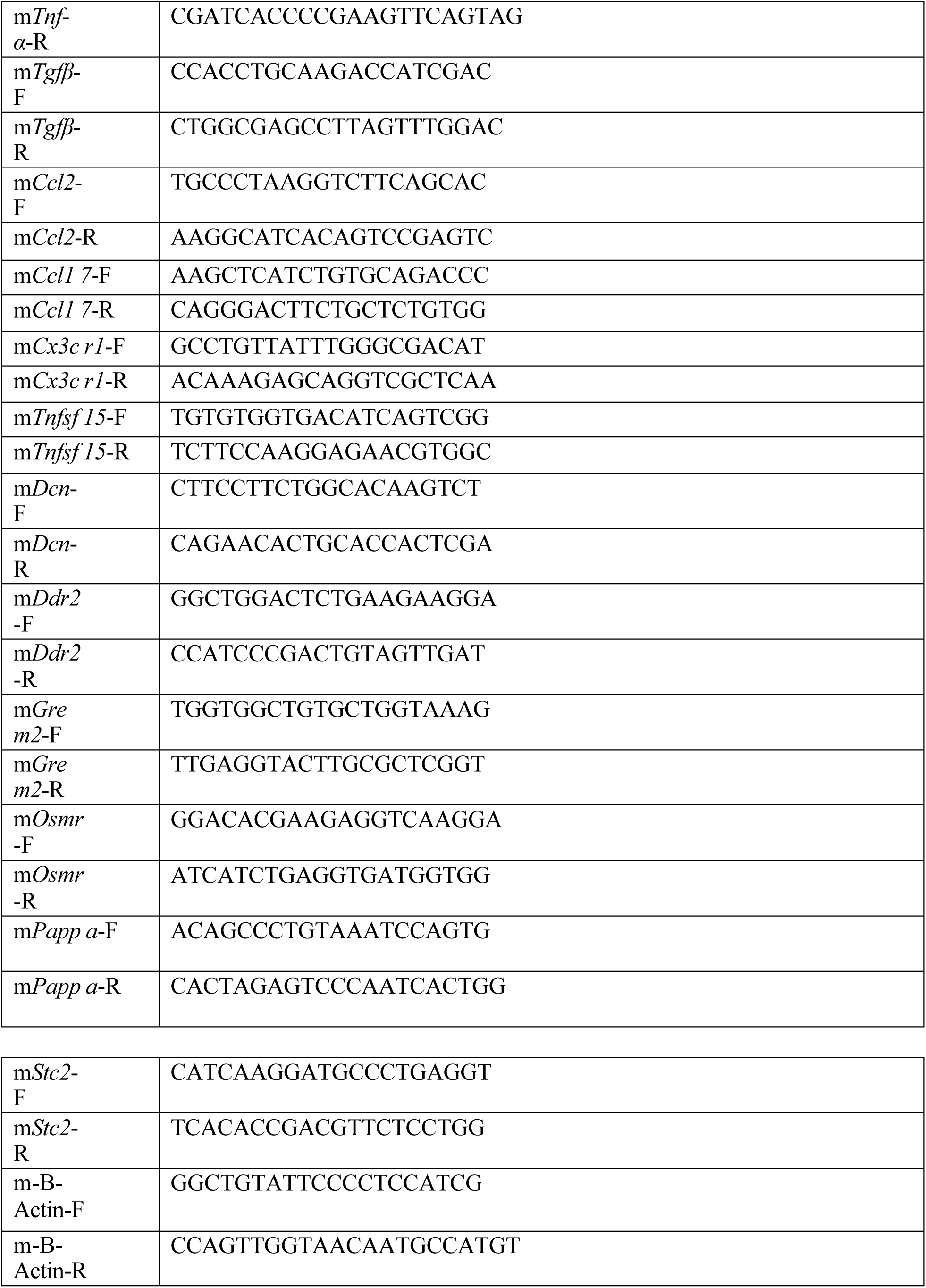
List of primers used for mutagenesis and qRT-PCR.

