## Supplemental Data (Figures and Tables) for "Pyridoxine supplementation confers protection against *SGPL1^R222Q^* variant sphingosine phosphate lyase insufficiency syndrome"

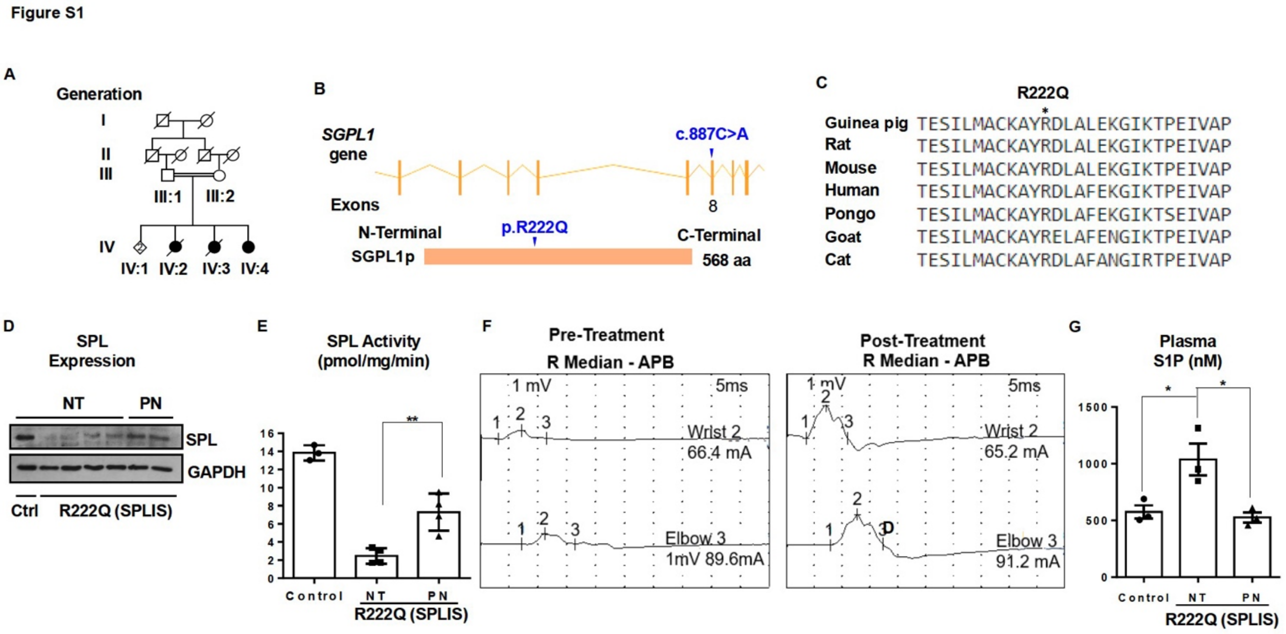


**Figure S1. Pyridoxine prevented disease progression in a patient with SPLIS-associated SPL$^{R222Q}$ variant.** (A) An extended consanguineous pedigree showed the one alive female patient (IV:4) with SPLIS-related symptoms. (B) *SGPL1* missense variant (c.887C>A) in exon 8 was supposed to cause a change in protein folding, resulting in an unstable SPL protein. (C) Sequence alignment of the region containing part of SPL amino acid sequence in guinea pig, rat, mouse, human, pongo, goat, and cat showing highly conserved nature of arginine. (D) SPL expression was measured by IB of whole cell extracts of fibroblasts from a healthy control (Ctrl) propagated for 24h in medium with no added pyridoxine, i.e., no treatment (NT), versus fibroblasts from a patient homozygous for the SPL$^{R222Q}$ variant (R222Q SPLIS) propagated in medium with no added PN for 24h followed by no further treatment (NT) or followed by propagation in medium containing 100 μM PN for 72h. GAPDH is a loading control. (E)

Treatment of $SPL^{R222Q}$ variant skin fibroblasts with pyridoxine increased SPL activity to ~25% of control fibroblasts. Untreated vs PN, $p < 0.005$. (F) Improvement in motor nerve conduction of the patient (IV:4) in post-B6 treatment. (G) Plasma S1P in the patient (IV:4) is reduced after two months of PN treatment. Aged match untreated vs post-B6 plasma S1P, $p < 0.04$.

Figure S2

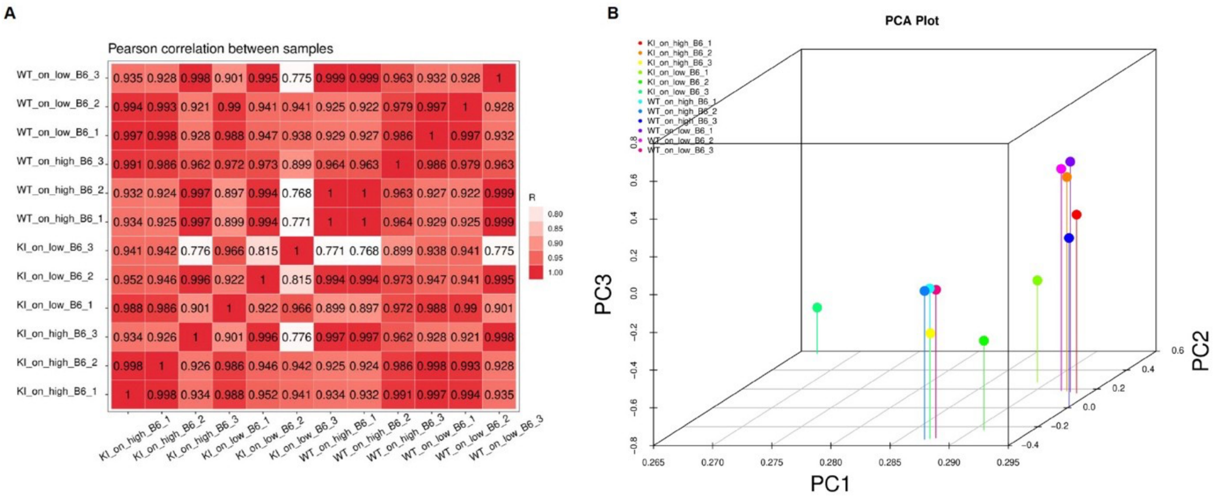


**Figure S2. PCA analysis and sample correlation.** (A) Pearson correlation is used to assess the linear relationship and similarity between different samples. (B) Principal component analysis (PCA) was used to simplify complex gene expression data, identify patterns, and visualize relationships between samples.

Figure S3

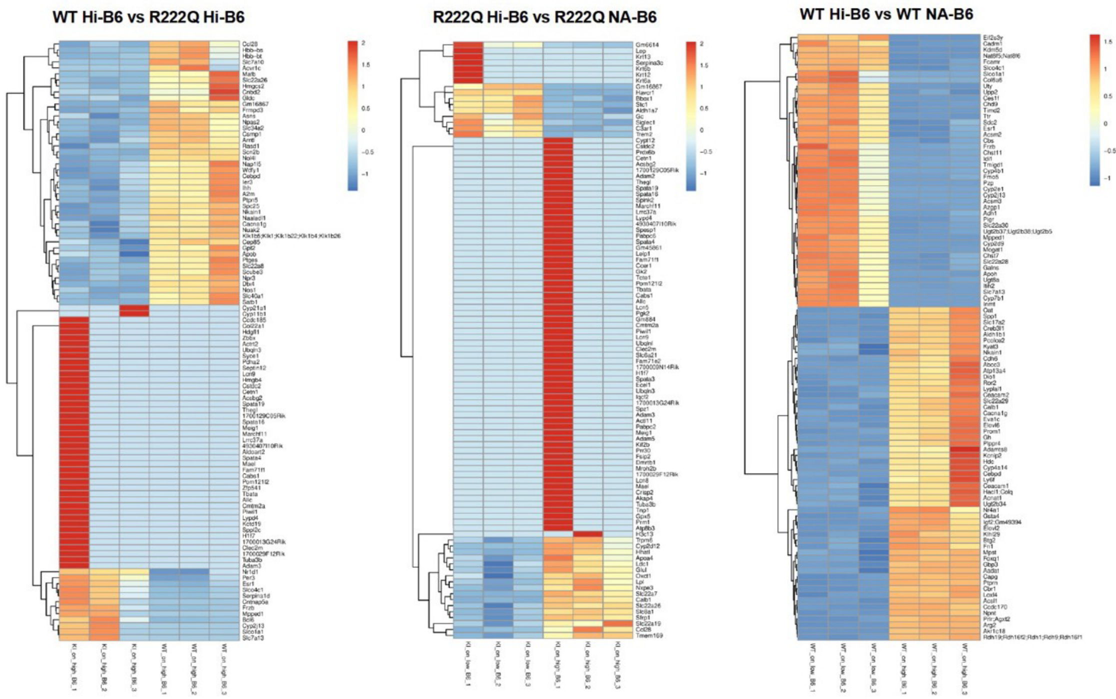


**Figure S3. Heatmap analysis of WT on Hi-B6, WT on NA-B6, SPL$^{R222Q}$ on Hi-B6 chow.** We excluded those genes that were common between the WT Hi-B6 vs the WT NA-B6 group to avoid the effect of low B6.

Figure S4

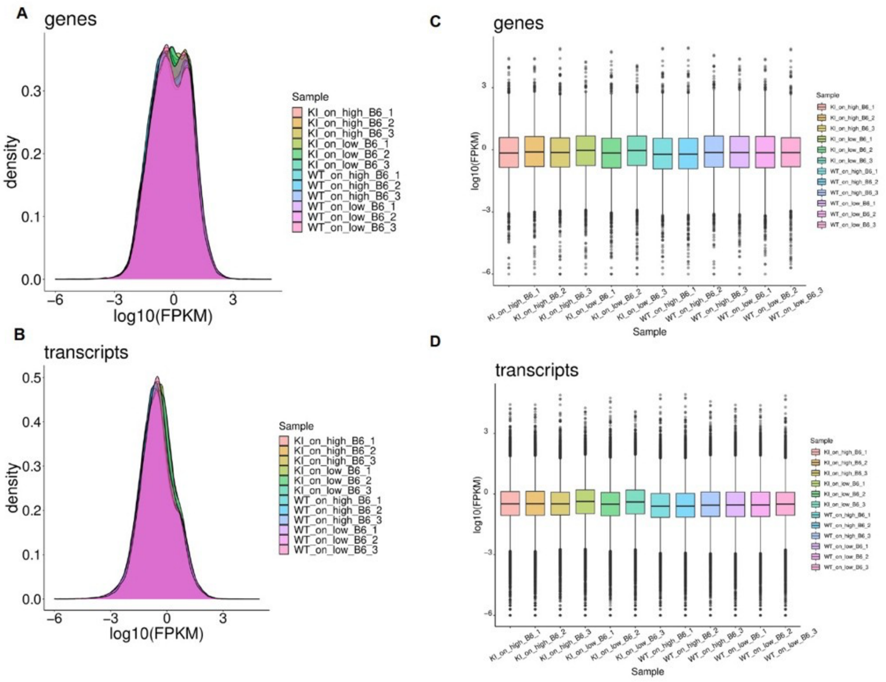


**Figure S4. Overview analysis of whole kidney transcriptome RNA-seq**. (A-D) Gene and transcript density levels (A&B) and boxplot expression (C&D) in different groups of samples were estimated by StringTie using FPKM (total_exon_fragments/mapped_reads(millions)×exon_length(kB)]).

Figure S5

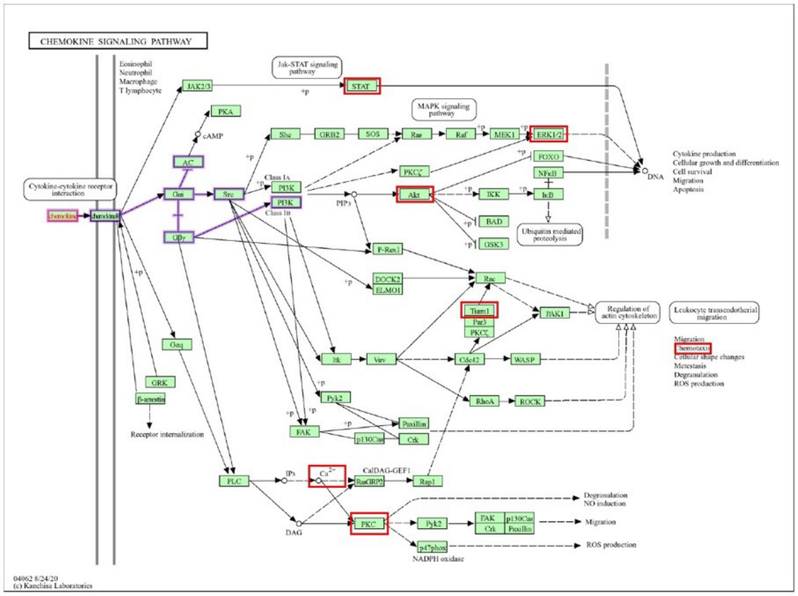


**Figure S5. KEGG pathway showing the involvement of cytokines in activating stress signals**
